# Mechanisms regulating reactivation pathways of *Toxoplasma gondii* as revealed by bradyzoite transgenesis

**DOI:** 10.1101/2022.07.21.500998

**Authors:** Sandeep Srivastava, David D. Hong, Krista Brooks, Amber L. Goerner, Edward A. Vizcarra, Emma H. Wilson, Michael W. White

## Abstract

Current approaches to find therapeutic solutions to treat and prevent reactivation of toxoplasmosis have suffered from limited accessibility to the relevant *Toxoplasma* stages and a lack of accurate in vitro developmental models. The loss of developmental competency in vitro that is exacerbated during the generation of transgenic tachyzoites is also a major impediment to understanding the molecular basis of bradyzoite recrudescence, which is the central parasite stage of reactivation. We have successfully modified *Toxoplasma* genes in the bradyzoite tissue cyst stage using ex vivo bradyzoite-based methods. Our new protocols validated the clonality of individual cysts and confirmed that single tissue cysts can robustly infect mice. We demonstrated these protocols by generating in vivo bradyzoites carrying a knockout of the *Toxoplasma* HXGPRT gene or the gene encoding the ApiAP2 transcription factor, AP2IX-9. Unexpectedly, the knockout of the AP2IX-9 gene in the Type II ME49EW strain eliminated one of the three developmental pathways initiated by the bradyzoite: host-dependent bradyzoite-to-bradyzoite replication. Our genetic protocols were further validated by producing in vivo bradyzoites lacking the bradyzoite-specific cyclin, TgCYC5. Interestingly, genetic ablation of TgCYC5 led to a large number of small cysts that formed from single mother cysts in mouse brain. Further study revealed the cause of the small-cyst phenotype in TgCYC5 knockout parasites was a disruption in the normal balance of bradyzoite subtypes that enhanced the bradyzoite-to-bradyzoite developmental pathway. These new data demonstrate the feasibility of generating transgenic parasites in a developmentally competent strain using ex vivo bradyzoite-based methods. Furthermore, by targeting the AP2IX-9 and TgCYC5 genes, whose transcripts are upregulated specifically in bradyzoites, we have clarified how these factors influence the pathways of tissue cyst recrudescence.

## Introduction

*Toxoplasma gondii* infects a wide range of nucleated cells in warm-blooded hosts but demonstrates a pronounced preference for immune-privileged tissues with limited immune surveillance, such as the central nervous system, eyes, and placenta [1]. Infected brain and muscle tissues serve as reservoirs for latent tissue cysts resulting in life-long infections [2]. Notably, toxoplasmosis is the most common opportunistic infection in advanced AIDS patients with low CD4 cell counts, necessitating effective prophylaxis [3]. In pregnant women lacking pre-existing antibodies against *T. gondii*, the parasite can cause acute, life-threatening infections in developing fetuses, leading to congenital toxoplasmosis. Unfortunately, no approved treatment effectively eliminates tissue cysts residing in the brain, posing a significant risk of recurrent reactivation in both symptomatic and asymptomatic individuals [4–8]. There is also concern about a drug-resistant “persistent population” of parasites that may survive after the clearance of the majority of the infection [9–11]. Thus, cyst maintenance, propagation and recrudescence are essential parts of the *Toxoplasma* life cycle responsible for disease and transmission.

Despite the importance of *Toxoplasma* reactivation to disease, the molecular mechanisms governing tissue cyst recrudescence remain poorly understood. The absence of an accurate model for bradyzoite developmental biology continues to impede advancements in understanding the parasite mechanisms responsible for chronic toxoplasmosis. Current *in vitro* laboratory-adapted protocols are limited to studying the early stages of tachyzoite-to-bradyzoite conversion and genetic methods that apply extensive serial cell culture passage in particular exacerbate this problem. *Toxoplasma gondii* boasts one of the most advanced genetic toolkits in eukaryotic research. They have provided critical insights into the tachyzoite cell cycle, intracellular invasion, and parasite-host cell interactions. However, these genetic methods are less effective for investigating the stages responsible for chronic infections and disease recrudescence due to the unique biology of bradyzoites. Nonetheless, several bradyzoite regulators, including BFD1 and ROCY1, have been identified alongside classical AP2 transcription factors. Both BFD1 and ROCY1 are essential for regulating tachyzoite conversion to bradyzoites [12, 13]. In this study, we introduce ex vivo bradyzoite protocols that address the limitations of current genetic approaches for bradyzoite research [14]. We present scalable methods for producing tissue cysts from animal models and describe novel genetic strategies that utilize ex vivo bradyzoites to generate transgenic *in vivo* tissue cysts. Using our newly developed protocols, we successfully generated a knockout of the bradyzoite-specific transcription factor TgAP2IX-9 and the bradyzoite-specific cyclin, TgCYC5. These knockouts had opposite effects on bradyzoite recrudescence leading to unique in vivo phenotypes in chronically infected mice.

## Results

### Maintenance and scale production of developmentally competent tissue cysts

The ME49 strain used in these studies (designated ME49EW) has been exclusively maintained for >20 years in vivo by alternating passage through resistant and sensitive mouse strains in order to sustain efficient tissue cyst production (Fig. 1A). The two mouse strains used in these studies, are the disease sensitive CBA/j strain and relatively disease resistant Swiss Webster (SWR/j) strain. From an infection of 10 cysts, the ME49EW strain consistently produced thousands of tissue cysts in CBA/j mice at 40-day post-infection (d.p.i.) and lower, but still robust cyst numbers in SWR/j mice (Fig. 1A). Alternating passage between CBA/j and SWR/j mouse strains was required to maintain robust tissue cyst numbers in mouse brain tissue (Fig. 1B). Repeated passage in CBA/j mice led to reductions in brain tissue cyst number that could be restored by a single passage through SWR/j mice (Fig. 1B, 30 d.p.i. cyst counts). The molecular basis of restoring declining tissue cyst numbers by passage in SWR/j mice is under investigation in our laboratories.

**Figure 1.**
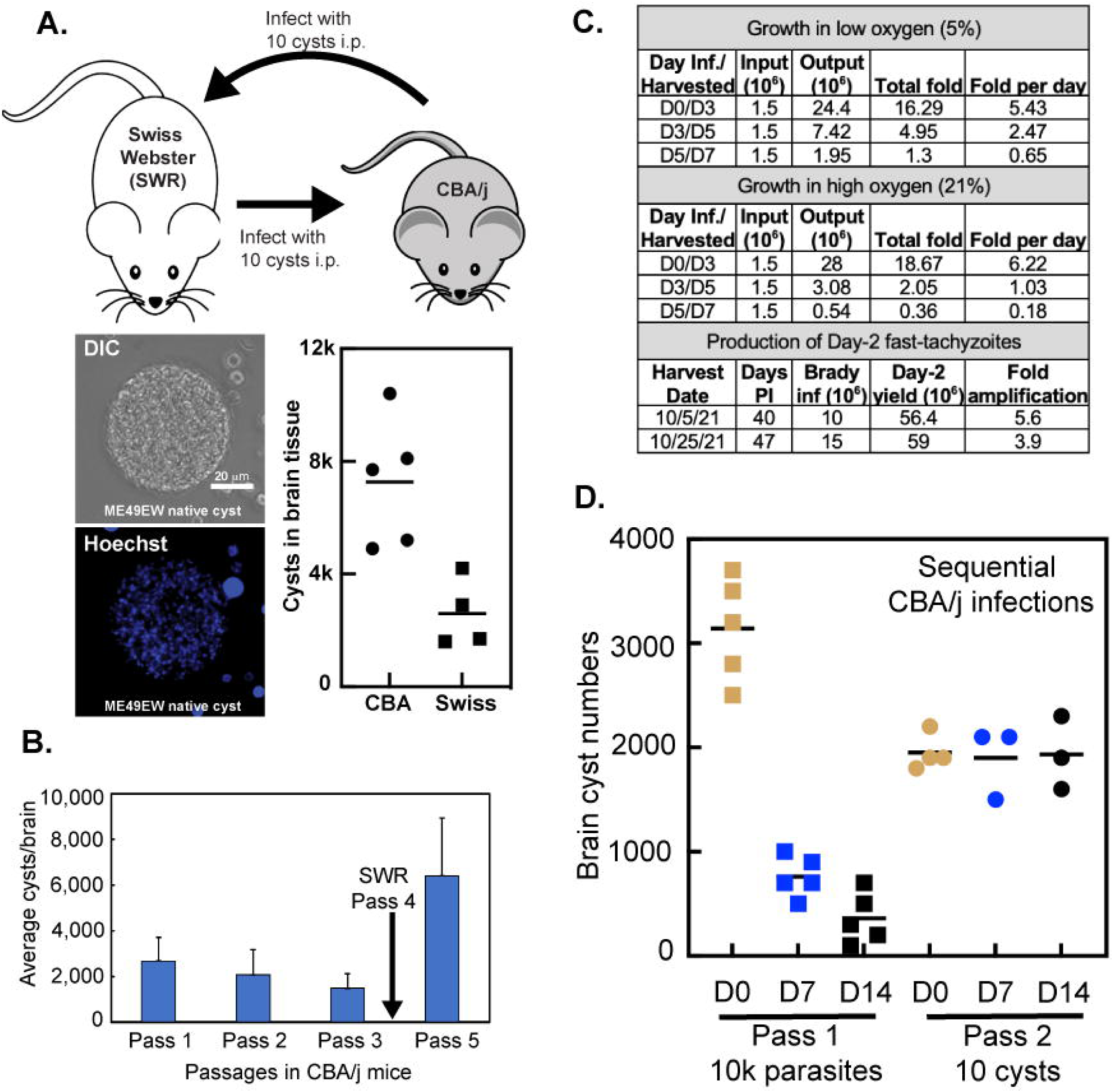
Alternating mouse strain infections stabilizes tissue cyst yields in brain cortex tissue. ME49EW strain is maintained in the relatively resistant Swiss Webster (SWR/j) mouse strain by a 10-cyst inoculation (i.p.) every 30-60 days. Tissue cysts from SWR/j brain homogenates are used to amplify tissue cysts numbers in the sensitive CBA/j mouse strain (10 cyst/mouse, 30-50 d.p.i.). **A.** Representative example of a ME49EW brain tissue cyst purified from mouse brain cortex tissue at 40 d.p.i. Brain tissue cyst numbers (cortex only) are higher in CBA/j versus SWR/j mice. **B.** Sequential passage in CBA/j mice can result in a progressive decline in tissue cyst number after multiple passages. If the single mouse strain passages are not excessive, a single passage through SWR/j mice can restore higher cyst numbers in the next CBA/j infection. Average cyst numbers for each CBA/j passage involved >5 mice/passage. **C.** ME49EW bradyzoite recrudescence infections in primary mouse astrocytes are influenced by oxygen gas concentrations. Parasite population growth is improved by lowering the oxygen levels to 5% as compared to standard atmospheric oxygen (21%) conditions in astrocyte cultures (D3/D5 and D5/D7) beyond the first astrocyte monolayer infected with excysted-bradyzoites (D0/D3). Note parasite replication in the first 24 h of bradyzoite-infected cultures is minimal (∼1.4 fold at D1). Relevant to nucleofection experiments, a 4-5-fold amplification of parasite numbers in Day-2 populations as compared to bradyzoite inoculations at Day-0 was reproducible (bradyzoites were obtained from mice ≥40 days post-infection). A reproducible shift to significantly slower parasite growth occurs beyond Day-5 in these cultures. **D.** Infections of CBA/j mice with Day-7 or Day-14 parasites from SWR/j ex vivo bradyzoite infections of astrocytes produce fewer brain cysts (blue and black filled squares) as compared to excysted-bradyzoite infections (orange filled squares). First round of CBA/j infections (Pass 1)=10,000 parasites i.p. A second round of CBA/j infections (Pass 2, 10 cysts i.p) using tissue cysts from Pass 1 mice (orange, blue or black filled squares) is shown. Average brain cortex tissue cyst numbers were determined from groups of five CBA/j mice at 30 d.p.i.

In the absence of a suitable laboratory model to investigate native bradyzoite biology, we optimized ME49EW tissue cyst and bradyzoite purification protocols to reliably yield an average of 1.4 million purified bradyzoites per CBA/j mouse (Fig. S1A and Protocol S1). The ME49EW/CBA/j model for the production of tissue cysts has been run at increased scale (25-50 mice/month) in our laboratories for several years and this model is reliable for obtaining native bradyzoites. Two key changes were introduced to the cyst isolation protocol that improved consistency and yields of tissue cyst purifications (see Protocol S1). **1.)** Only the mouse cortex was harvested for tissue cyst processing. Anatomical mapping of mouse brain infections [15] recently determined the majority of tissue cysts are localized to the brain cortex and not in the cerebellum. In our protocols, removing the white fibrous cerebellum improved homogenization of brain tissue without significant loss of cysts (<10% losses). **2.)** Whole brain cortices were kept cold (0-4°C) in 1xPBS for ∼16 h. Cold incubation of mouse brain tissue improves homogenization, and cyst yields with no discernable effect on tissue cyst viability or the ability of excysted bradyzoites to invade host cells and initiate recrudescence. Homogenates of freshly harvested brain tissue suffered cyst losses (30-50% loss) during percoll gradient purification due to cyst trapping in incompletely homogenized tissue.

### Establishing ex vivo cell culture models

In order to take advantage of the ex vivo model of bradyzoite recrudescence [14] for the purpose of developing new genetic strategies, we used primary mouse neonatal astrocytes to expand of tachyzoite populations following ex vivo bradyzoite infection. Initial ME49EW bradyzoite infections of HFF cells yielded poor medium-term ME49EW parasite growth, and were therefore, not used for developing new genetic protocols [14]. We first evaluated ME49EW bradyzoite-initiated growth in astrocyte specialized media at two oxygen gas conditions (5% versus 21%). We reasoned oxygen conditions might influence ex vivo bradyzoite models due to the hypoxic oxygen environment of brain tissue where tissue cysts reside [16]. Short term parasite growth (Day 1-3) in bradyzoite-infected astrocytes was not affected by oxygen levels, however, in astrocyte cultures beyond Day-3, parasite growth was negatively affected by atmospheric oxygen levels (21%) (Fig. 1C). A similar oxygen effect on ME49EW bradyzoite recrudescence was also observed in HFF cells [14]. As a consequence of these experiments, lower oxygen conditions (5%) in hypoxic chambers were used to cultivate ME49EW parasites throughout these studies.

It was important to evaluate whether cultivation of native ME49EW parasites in astrocytes would affect the capacity to form tissue cysts in mice. Based on the positive recovery of cyst numbers after passage through SWR/j mice (Fig. 1B), we evaluated using ex vivo bradyzoites from SWR/j mice as the starting source for transfection experiments where sequential passage through astrocytes (to obtain fast-growing tachyzoites for transfection) and mice will be required (see Protocol S2). Excysted bradyzoites from SWR/j mice were used to infect astrocytes and after 7 and 14 days in culture 10,000 parasites were used to infect mice. Cyst counts were then determined at 30 days (d.p.i.). As we previously determined for Day-7 parasites from astrocytes [14], direct infections with Day-14 parasites yielded lower cyst numbers compared to mouse infections with ex vivo bradyzoites (Fig. 1D). To determine if this reduction in cyst yield was recoverable, a second round of infections using 10 tissue cysts i.p. (standard dose) obtained from mice from the the first-round of infections demonstrated similar cyst counts independent of the original source (day 0, 7 or 14) of infection indicating cyst formation is recoverable when ex vivo ME49EW bradyzoites were isolated from SWR/j mice. By contrast, ME49EW bradyzoites isolated from CBA/j mice and then cultured in astrocytes for 14 days did not show a similar tissue cyst recovery in secondary mouse infections (Fig. S1B). For the purposes of genetically modifying native parasites, ex vivo bradyzoites from SWR/j mice were the preferred source for initiating transfection protocols.

### Engineering of transgenic tissue cysts

Current genetic protocols rely on highly modified tachyzoite strains (e.g. Δhxgprt, Δku80, TIR1- and GFP-expression) that were well adapted to grow in HFF cells, and as a consequence poorly form tissue cysts in mice when compared to native strains like ME49EW [14]. Our goal was to develop an approach for genetically modifying *Toxoplasma*, while preserving developmental competency. The new method was developed and tested using a plasmid design that targeted the HXGPRT gene for replacement with a DNA insert containing the pyrimethamine-resistant TgDHFR selectable marker and GFP and firefly luciferase reporters (see Fig. 2A). The key to a development-sparing genetic model was the fast-replicating tachyzoite stage that forms early in ME49EW ex vivo bradyzoite recrudescence [14]. The rapid growth of these parasites provides ∼5-fold parasite amplification by Day 2 (Fig. 1C) that together with the reduced number of parasites needed (1×10^7^) for nucleofection necessitated fewer infected mice in these protocols (5-8 mice per nucleofection). Robust growth of Day-2 ME49EW parasites was the key ingredient for comparatively high transformation frequency (0.58%, Fig. 2B) equal to or exceeding the Type-I, RH tachyzoite (0.44%, Fig. S1C) and ∼10-fold higher than HFF-adapted ME49B7 tachyzoites (Fig. S1C). Conventional plaque assays to measure transformation were not possible in the ex vivo bradyzoite recrudescence model, and thus, transformants were estimated by flow cytometric quantification of GFP+ parasites following three days of selection in 0.5 μM pyrimethamine (Day 3 through Day 6 post-bradyzoite infection).

**Figure 2.**
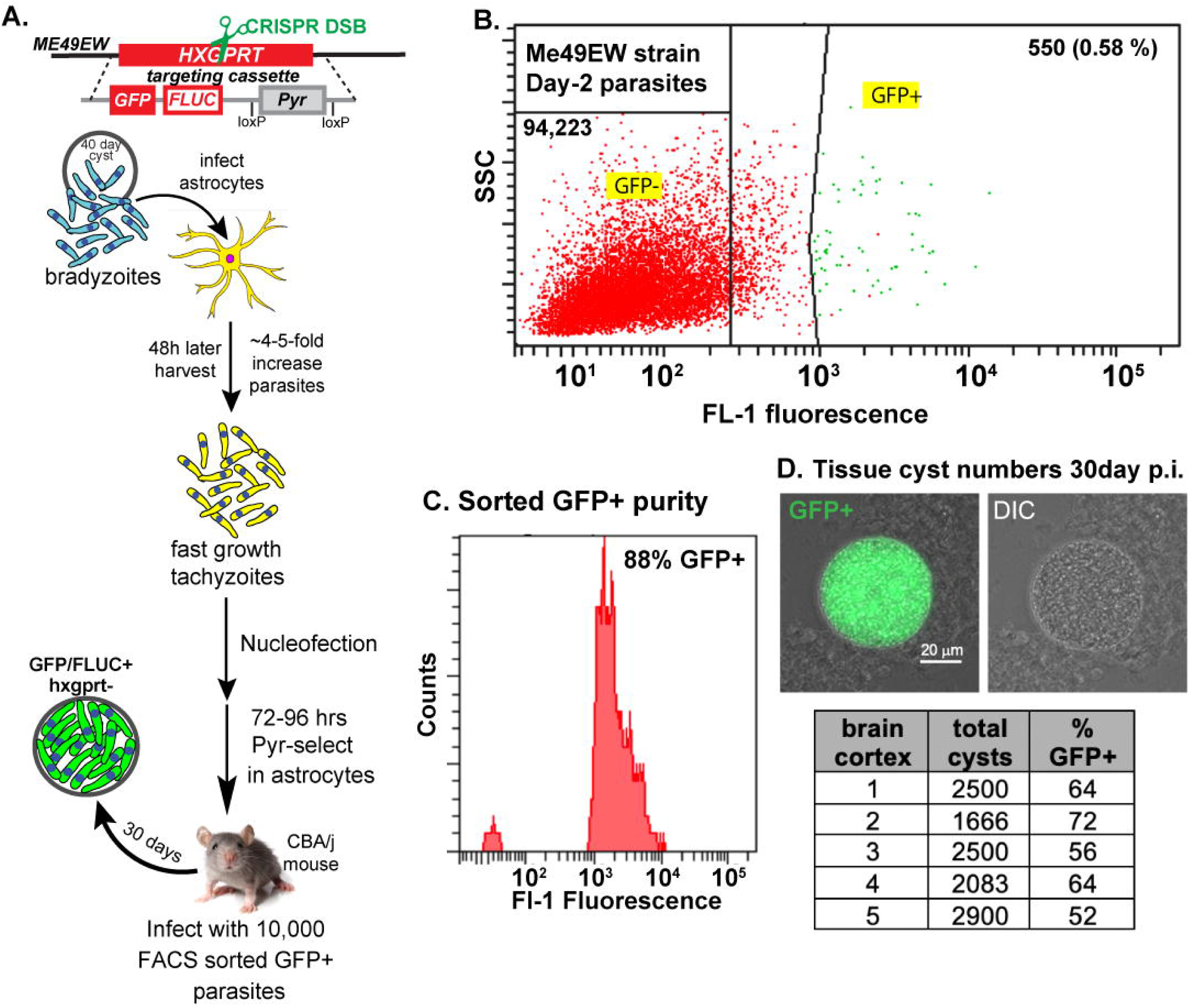
Knockout of the HXGPRT gene in ME49EW bradyzoites. **A.** Overview of the protocol for generating transgenic tissue cysts in mice. For complete methods see Protocol S2. **B.** The HXGPRT gene was targeted for CRISPR-assisted knockout in ME49EW Day-2 recrudescent parasites. Following nucleofection and 72 h post-pyrimethamine (0.5μM) selection in astrocytes, GFP+ parasites were FACS sorted (55,000 total per sort). Gates defining GFP-versus GFP+ parasites are shown. Note that GFP+ represent 0.58% of total parasites analyzed by flow cytometry. Comparable nucleofection and FACS sorts were performed for Type 1 RH strain and ME49B7 tachyzoites (see Fig. S1B) **C.** The sorted GFP+ ME49EW parasites reanalyzed by flow cytometry were 88% GFP+ representing a 150-fold enrichment over the pre-sort population. **D.** GFP+ sorted parasites were used to infect five CBA/j mice (10,000 parasites i.p./mouse). At 30 d.p.i., brain cortex tissue was harvested and total tissue cysts and GFP+ expression determined. A representative image of GFP+ transgenic tissue cyst is shown (live GFP fluorescence compared to DIC image, 1000x).

To limit the cell culture of ME49EW recrudescing parasites, GFP+ transformants were FACS sorted at Day-7 post-bradyzoite infection (3 days of pyrimethamine selection) and >50,000 GFP+ parasites were collected. We reanalyzed sorted parasites by flow cytometry and confirmed a >150-fold GFP+ enrichment over pre-sorted populations (88% GFP+, Fig. 2C). A group of five CBA/j mice were infected using a dose of 10,000 GFP+ parasites i.p. per mouse. At 30 d.p.i., cysts were harvested, and total cyst counts and percent GFP+ determined microscopically in brain homogenates. Total cyst counts reflected a burden similar to that of ME49EW parent infections with an average of 2,329 tissue cysts per brain. We expected both GFP+ and GFP-cysts as the FACS sort was not 100% GFP+ parasites (Fig. 2C), and thus, the number of GFP+ tissue cysts in mouse brain varied from 52% to 72%. Notably, tissue cysts harbored either 100% GFP+ or 100% GFP-bradyzoites, no mixed tissue cysts were observed (Fig. 2D). The next step in the goal to produce transgenic tissue cysts was to obtain clonal GFP+ cyst isolates.

### Cloning and preserving ME49EW transgenic tissue cysts

The most common protocol for cloning transgenic tachyzoites utilizes limiting dilution of parasites in 96 well plates. Unfortunately, this approach requires extensive cell culture risking the rapid loss of developmental competency. The tachyzoite parasitophorus vacuole is formed from a single parasite infection, and thus, each vacuole harbors clonal daughter parasites. Tissue cysts are derived from individual vacuoles [17], and are therefore, also expected to be clonal. Our GFP cyst data supports this, as all bradyzoites within a cyst are of a single GFP phenotype either positive or negative and no cysts were observed containing bradyzoites that were positive and negative for GFP. This opened the possibility that cloning transgenic isolates by single tissue cyst infections in mice was feasible. We first evaluated the reliability of very low dose ME49EW cyst infections in mice by mixing GFP+ and GFP-cysts 1:1 and infecting CBA/j mice with 2 cysts. At 30 d.p.i., tissue cyst numbers and the distribution of GFP+ versus GFP-parasites within tissue cysts were determined (Fig. 3A). There was a 1 in 4 theoretical chance that a 2-cyst infection of mice would be homogeneous GFP+ or GFP- (25% each). All 20 mice were successfully infected (one mouse died from the *Toxoplasma* infection before 30 d.p.i.) with 5 mice having 100% GFP+ brain cysts and 3 mice having 100% GFP-cysts. The remaining 11 mice were infected with various combinations of 100% GFP+ and 100% GFP-tissue cysts in ratios of ∼1:3, ∼1:4, ∼1:5, and some ∼1:20. Because we did not control for cyst size (i.e. variable bradyzoite numbers) these types of mixtures were expected. Importantly, there were no individual cysts harboring mixtures of GFP+/GFP-parasites in the hundreds of cysts we examined microscopically confirming the clonality of bradyzoites within individual cysts. The infection success with low cyst numbers led us to test methods to isolate single cysts in brain homogenates or from percoll-purified cysts. Limiting dilution of cysts in 96 well optical plates (1 cyst/100μl) followed by microscopic screening was an effective strategy to isolate single cysts (see Protocol S3 for details). The contents of wells with single cysts were loaded into syringes and used to infect CBA/j mice (percoll or homogenate diluted preparations). All mice were successfully infected and yielded hundreds to thousands of clonal tissue cysts in brain tissue at 30 d.p.i. (Fig. 3B).

**Figure 3.**
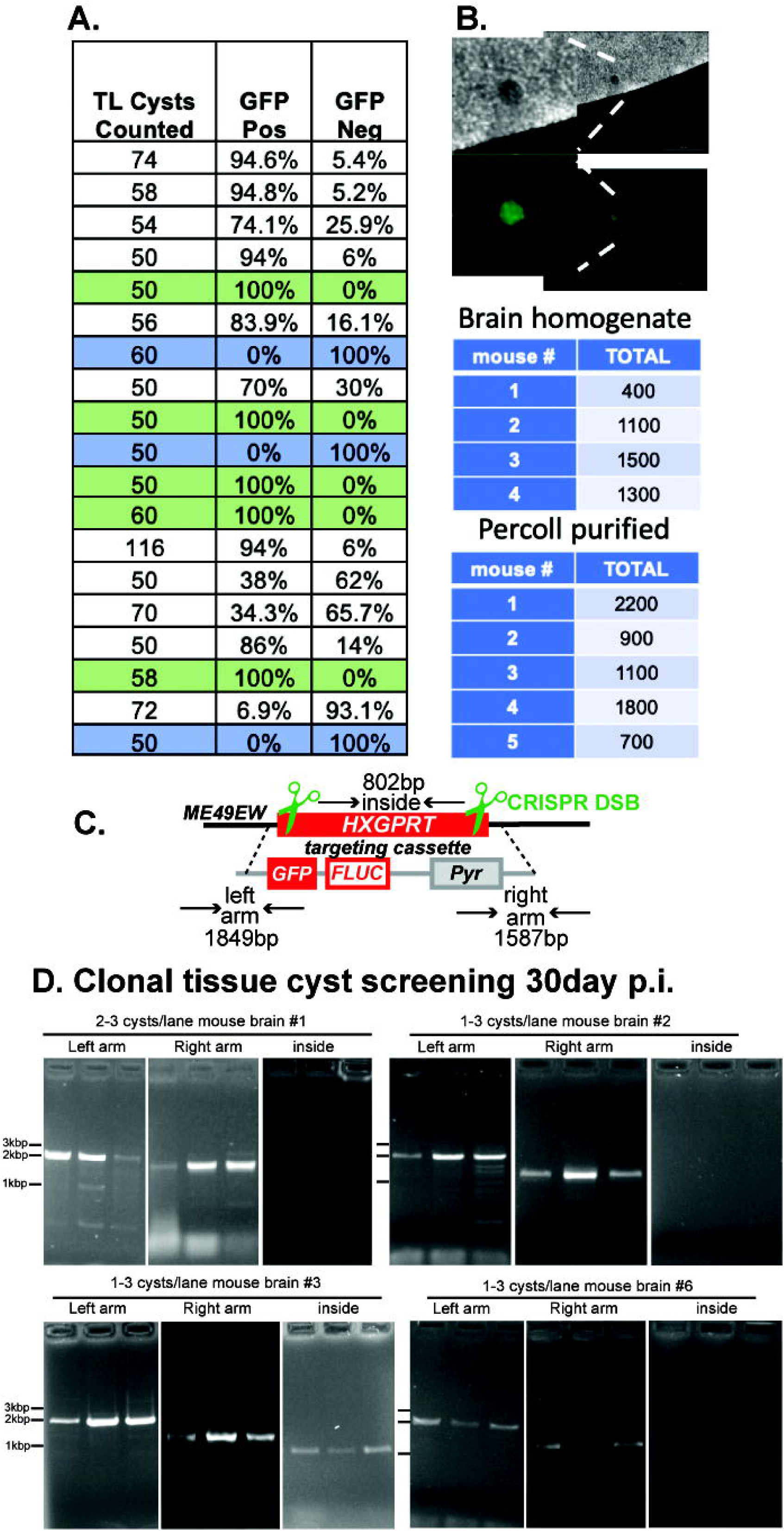
Cloning transgenic tissue cysts. **A.** GFP+ tissue cysts were mixed 1:1 with GFP-tissue cysts (ME49EW strains) and diluted to 2 cysts/0.20 ml of PBS. Twenty CBA/j mice were infected i.p with 2 cysts/mouse (one mouse died before cyst analysis). At 30 d.p.i., cortex brain tissue was harvested from each mouse and tissue cysts purified by percoll gradient (Protocol S1). A minimum of 50 live tissue cysts from each mouse were evaluated for GFP+ expression by fluorescence microscopy. **B.** Single tissue cysts are infectious and produce substantial tissue cyst numbers in CBA/j mice. Single GFP+ tissue cysts from brain homogenates or following percoll gradient purification were isolated by limiting dilution in 96 well optical plates. At 30 d.p.i., tissue cyst counts in brain homogenates from the nine infected mice indicated were determined. Note, no GFP-negative tissue cysts were observed in mice infected with a single GFP+ cysts (all 100% GFP+). **C.** Overview of HXGPRT knockout strategy and PCR screening strategy. The *Toxoplasma* HXGPRT coding region was replaced by a three gene cassette of GFP, firefly luciferase, and pyrimethamine-resistant DHFR. **D.** PCR analysis of the knockout of the TgHXGPRT gene in ME49EW tissue cysts was confirmed in single cyst infections of mice. The presence of the correct left and right arm DNA fragments in ME49EW::*Δhxgprt* parasite gDNA verified the correct double cross-over had occurred in the clone (left/right arm gels). The failure to amplify a TgHXGPRT 802bp internal gDNA coding fragment (inside gels) indicated a clean TgHXGPRT knockout.

To determine if the GFP+ phenotype was stable, we cloned HXGPRT knockout parasites (Fig. 3C and D) using cysts from a second nucleofection, followed by a GFP+ FACS sort and CBA/j infection. At 30 d.p.i., 4 mice were 100% GFP+ and one mouse was 33% GFP+. Tissue cysts in brain homogenate from one mouse harboring GFP+ cysts was diluted to 1 cyst/100 μl in PBS and plated into 96 well optical plates. Eight wells that appeared to contain single GFP+ cysts were identified and the total PBS volume of each well used to infect a CBA/j mouse (i.p.). At 30 d.p.i., brain cortex tissue was harvested, and cyst counts and GFP expression determined. One mouse had no cysts, while another only had GFP-cysts indicating a contaminating small cyst may have been missed during screening. The remaining six mice had GFP+ cyst numbers ranging from 100 to 8,000 cysts per brain with no GFP-cyst contamination. We selected four homogenates to move forward to PCR screens: Brain #1-600 cysts, Brain #2-1,600 cysts, Brain #3-700 cysts, and Brain #6-8,000 cysts. The brain homogenates were diluted to 1 cyst/100 μl with PBS and plated in 96 well optical plates. DNA was purified from selected wells containing 1-3 cysts and analyzed using PCR (Fig. 3C and D). All DNA templates produced the right and left arm DNA fragments indicating a successful HXGPRT gene knockout. Homogenates #1, #2, and #8 cysts did not amplify an internal DNA fragment from the native HXGPRT gene indicating they were clean HXGPRT knockouts, while brain #3 DNA template did amplify the internal fragment indicating this mouse was likely dually infected with the HXGPRT knockout and ME49EW parent parasites. Homogenates containing cysts from mouse brain #1, #2, and #8 were frozen at -80°C using Protocol S4.

### Engineering an AP2IX-9 gene knockout in ME49EW tissue cysts

A major class of transcription factors in *Toxoplasma* are nuclear proteins distantly related to the APETLA family of transcription factors of plants [18]. The *Toxoplasma* genome encodes >60 of these factors, one of the largest collections of ApiAP2 genes in the Apicomplexa family of parasites. A subset of ApiAP2 factors are developmentally expressed and have functions in regulating *Toxoplasma* life cycles [19–23]. Recently, we determined that AP2IX-9 was the highest expressed ApiAP2 mRNA in ME49EW bradyzoites from chronically infected mice [14]. This result was unexpected as previous studies only detected transient expression of AP2IX-9 mRNA when tachyzoites were induced to differentiate in vitro to bradyzoites under alkaline-media conditions [20, 21]. Because of this discrepancy, we re-examined the function of AP2IX-9 in ME49EW in vivo bradyzoites by knocking out the AP2IX-9 gene using the new ex vivo bradyzoite genetic methods (Fig. 2A). At 30 d.p.i., we isolated single GFP+ tissue cysts from brain homogenates and infected CBA/j mice and verified the AP2IX-9 knockout in this clonal round of infection using PCR (Fig. S2A). We also confirmed the absence of AP2IX-9 mRNA in this clone (Fig. S2A) and verified the gene deletion in chromosome IX (Fig. S2B). Groups of CBA/j mice were infected with ME49EW::Δ*ap2IX-9* or ME49EW parent strain tissue cysts (10 cysts i.p./mouse) and brain tissue cyst numbers quantified at 30 d.p.i. Knockout of AP2IX-9 in the ME49EW strain did not significantly diminish tissue cyst formation in CBA/j brain tissue (Fig. 4A, B) nor were there significant differences in cyst sizes for AP2IX-9KO or ME49EW bradyzoites (Fig. 4C).

**Figure 4.**
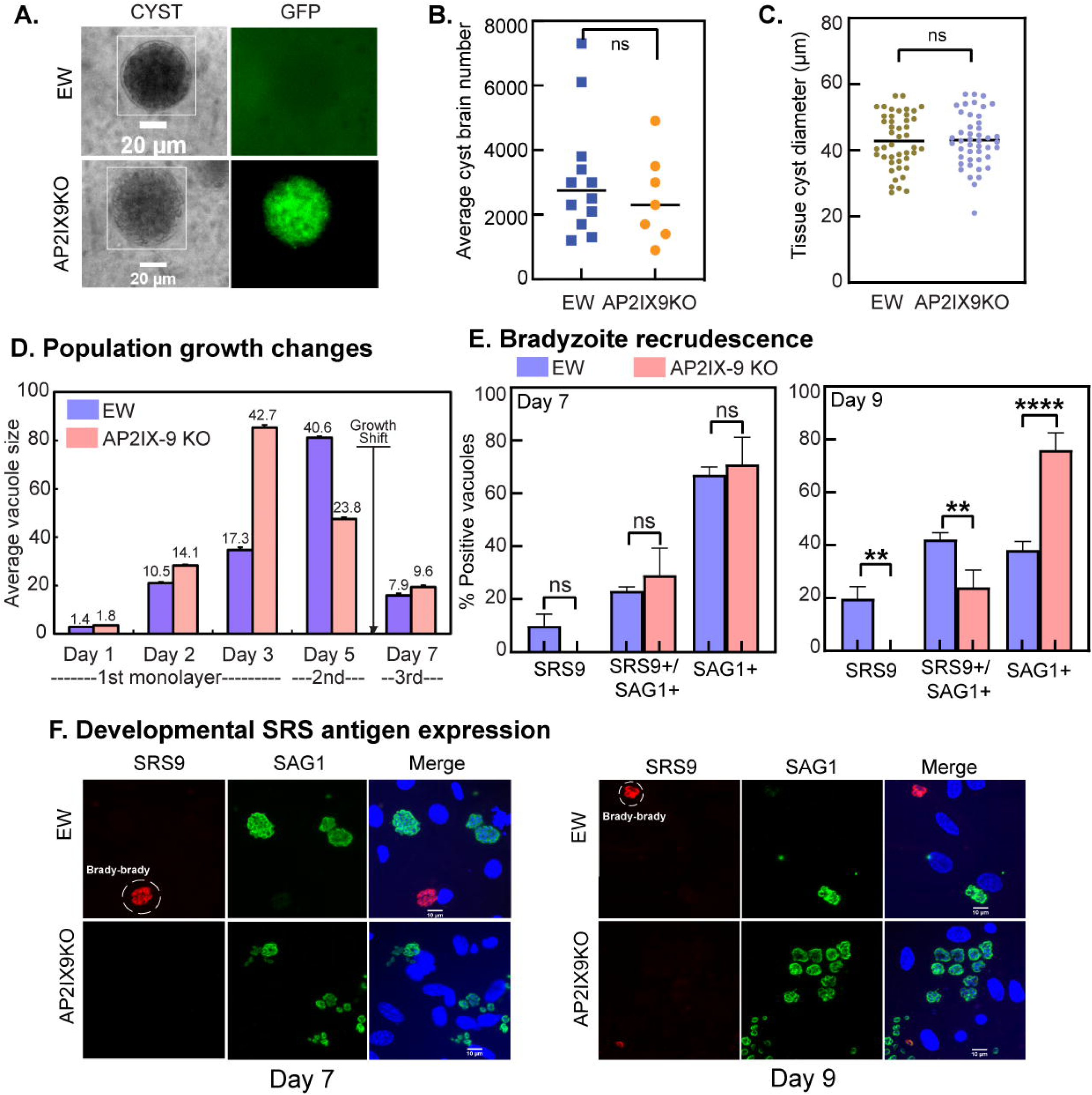
AP2IX-9 is required for bradyzoite-to-bradyzoite recrudescence. **A.** Representative ME49EW::Δ*ap2IX-9* tissue cyst in brain homogenate is GFP+, while ME49EW parent tissue cysts is GFP-. **B.** Quantification of tissue cysts in 40-day infections of CBA/j mice with ME49EW::Δ*ap2IX-9* or ME49EW parasites. **C**. Tissue cyst diameters were analyzed using ImageJ software. A total of 250 cysts were isolated and cloned into 96-well plates with microscopy images captured prior to measurement. No significant differences in tissue cyst size was observed in ME49EW::Δ*ap2IX-9* and ME49EW parent infections. **D.** Purified ME49EW::Δ*ap2IX-9* bradyzoites initiate the identical growth transitions (fast growth followed by switch to slow growth) in murine astrocytes as the parent ME49EW strain. **E. and F.** The timing of SRS9 and SAG1 surface antigen expression during ME49EW::Δ*ap2IX-9* bradyzoite recrudescence was quantified in four sequentially infected astrocyte monolayers. ME49EW::Δ*ap2IX-9* and ME49EW parent strains are designated as AP2IX-9KO and EW, respectively.

We next purified ME49EW::Δ*ap2IX-9* bradyzoites to further study the role of AP2IX-9 in ME49EW bradyzoite recrudescence. Primary mouse astrocytes were infected with 10,000 ME49EW::Δ*ap2IX-9* or ME49EW bradyzoites from in vivo cysts followed by the analysis of growth and development over 7-14 days. Previous analysis of growth kinetics initiated by ex vivo bradyzoite infections of astrocytes revealed a parasite-intrinsic 24 h awakening period followed by rapid parasite replication that uniformly slows between days 5-7. Furthermore, in addition to the classical recrudescence of bradyzoite-to-tachyzoite differentiation, there is host cell-dependent support of bradyzoite-bradyzoite replication representing ∼15% of the vacuoles at day 7 [14]. Analysis of ME49EW::Δ*ap2IX-9* parasites revealed that the parasite intrinsic growth pattern remains intact in the absence of AP2IX-9 (Fig. 4D). However, strikingly, bradyzoite-to-bradyzoite (SRS9+ only) replication seen in the ME49EW parent was absent in ME49EW::Δ*ap2IX-9*recrudescing populations (Fig. 4E, F). These results demonstrated that AP2IX-9 is required for direct bradyzoite-to-bradyzoite replication.

### Genetic knockout of TgCYC5 cyclin in ME49EW cysts

*Toxoplasma* cell cycle is faithfully regulated by checkpoint mechanisms involving cyclin machinery [24] with one cyclin, TgCYC5, uniquely expressed in the bradyzoite stage [25]. We investigated the in vivo function of TgCYC5 by applying our ex vivo bradyzoite-based genetic protocols to successfully knockout the TgCYC5 gene in ME49EW bradyzoites (Fig. S2A). Deletion of the TgCYC5 locus was confirmed by PCR of single GFP+ tissue cyst and the deletion of TgCYC5 gene in chromosome 1a (Fig. S2B). As with the AP2IX-9 knockout, the successful recovery of clonal cysts lacking the TgCYC5 gene in and of itself demonstrated that this protein was not essential for tissue cyst development. The phenotype of the AP2IX-9 knockout was manifest in bradyzoite recrudescence; we therefore explored this possibility for the ME49EW::Δ*cyc5* bradyzoites. Groups of CBA/j mice were infected with ME49EW::Δ*cyc5* or ME49EW parent, tissue cysts isolated 30 days post infection and free bradyzoites obtained by excystation. Primary murine astrocytes were infected with ME49EW::Δ*cyc5* or ME49EW parent bradyzoites and the progression of parasite recrudescence followed over a period of 9 days. ME49EW::Δ*cyc5* and ME49EW bradyzoites successfully invaded murine astrocytes and followed the same growth pattern for the first three days post-infection (Fig. 5A). Beyond Day-3, the growth of ME49EW::Δ*cyc5* parasites dramatically slowed as reflected in smaller vacuole sizes and population yields. The reduction in growth of ME49EW::Δ*cyc5* parasites was associated with a early re-expression of bradyzoite surface antigen SRS9 (while also expressing SAG1+) by Day 5 (Fig. 5B and C). Two days later (Day 7) the parent ME49EW parasites also slowed down growth and re-expressed bradyzoite-specific SRS9 indicating that TgCYC5 was required for extending the period of tachyzoite replication and preventing early shifts in tachyzoite-to-bradyzoite development.

**Figure 5.**
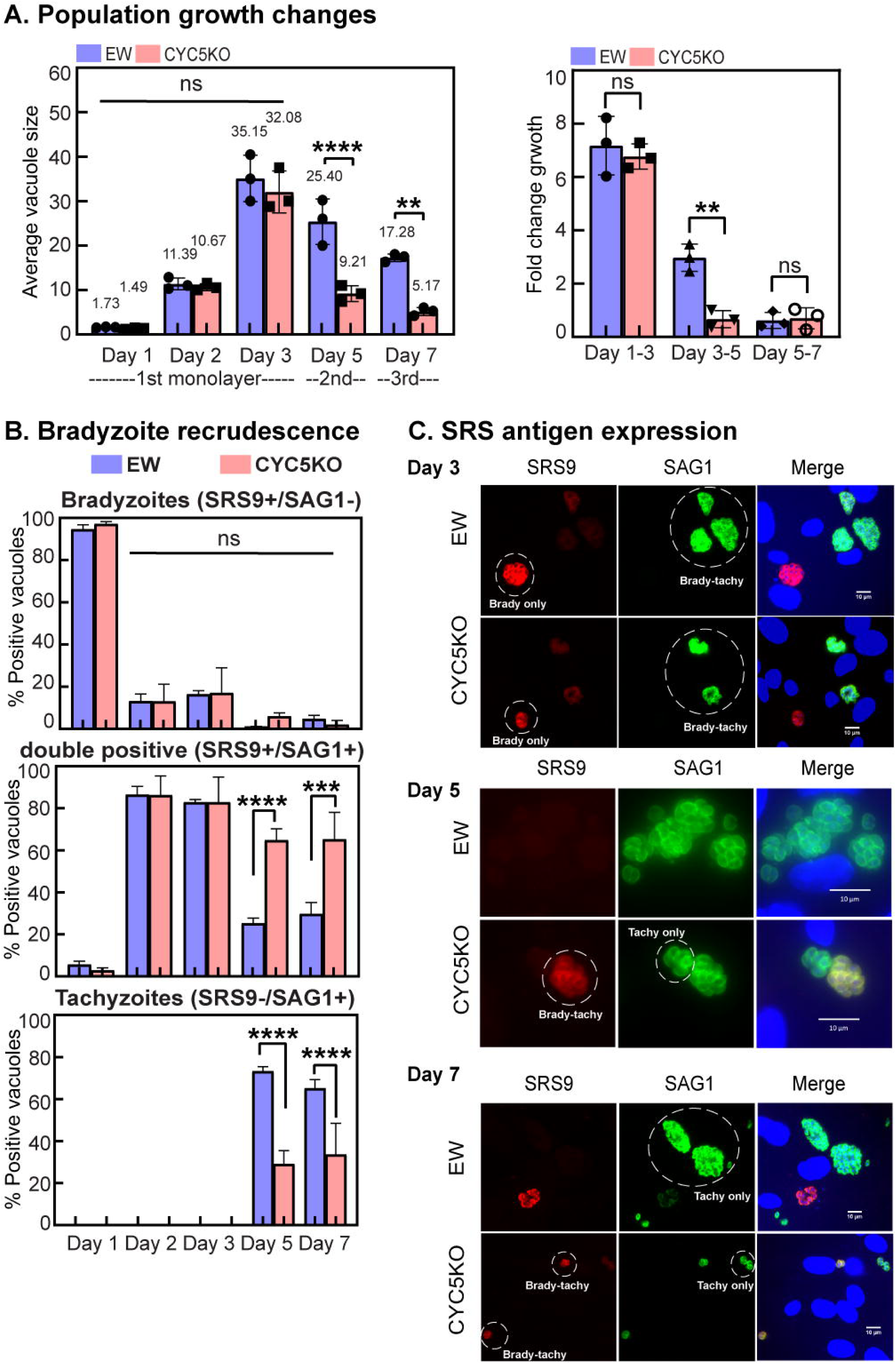
TgCYC5 regulates bradyzoite recrudescence in primary astrocytes. **A.** Purified ME49EW::Δ*cyc5* bradyzoites initiate identical reawakening (D0-D1) and early fast-growth periods (D2-D3) as the parent ME49EW strain. **B.** The timing of SRS9 and SAG1 surface antigen expression during ME49EW::Δ*cyc5* bradyzoite recrudescence compared to ME49EW parent strain was quantified in infected primary astrocytes. Note, the opposite pattern of SRS protein expression with ME49EW parasites converting primarily to tachzyoites (see bottom graph, SAG1+ only), whereas ME49EW::Δ*cyc5* parasites dramatically re-expressed SRS9+ (double positive, middle graph) in Day-5 infected astrocytes. **C.** Representative antigenic expression of SRS9 and SAG1 on days 3, 5, and 7 post-bradyzoite infection; SRS9+ is shown in red, while SAG1+ is show in green.

### TgCYC5 regulates daughter tissue cyst formation in vivo

While the knockout of TgCYC5 did not prevent tissue cyst formation, it did lead to visually smaller cysts compared to the ME49EW parent (Fig. 6A-C). The average ME49EW parent cyst was 35 µm, whereas ME49EW::Δ*cyc5* cysts were significantly smaller at 25 µm (Fig. 6B). Intriguingly, ME49EW::Δ*cyc5* bradyzoite infections of CBA/j mice yielded nearly 2.6-fold more cysts (19,700 average cysts/mouse brain) than the ME49EW parent (7,500 average cysts/mouse brain). In order to better understand why the TgCYC5 knockout led to increased numbers of small cysts, we conducted immunohistochemistry on brain slices from CBA/J mice infected with ten ME49EW::Δ*cyc5* or ME49EW parent cysts. Mice infected with the ME49EW::Δ*cyc5* strain developed small daughter cysts (∼35% cysts) adjacent to a large mother cyst, possibly sharing a common cyst wall, as observed by DBA staining (Fig. 6D). In addition, multiple small cyst clusters could be found adjacent to the mother cyst, suggesting recurrent and continuous daughter cyst formation might be occurring in ME49EW::Δ*cyc5* parasites compared to the EW parent strain (Fig. 6D). By contrast, most ME49EW parent cysts (85%) lacked evidence of daughter cyst formation. This in vivo phenotype of the ME49EW::Δ*cyc5* parasites is consistent with the early emergence of bradyzoites in the ex vivo recrudescence model associated with this mutant strain (Fig. 5).

**Figure 6.**
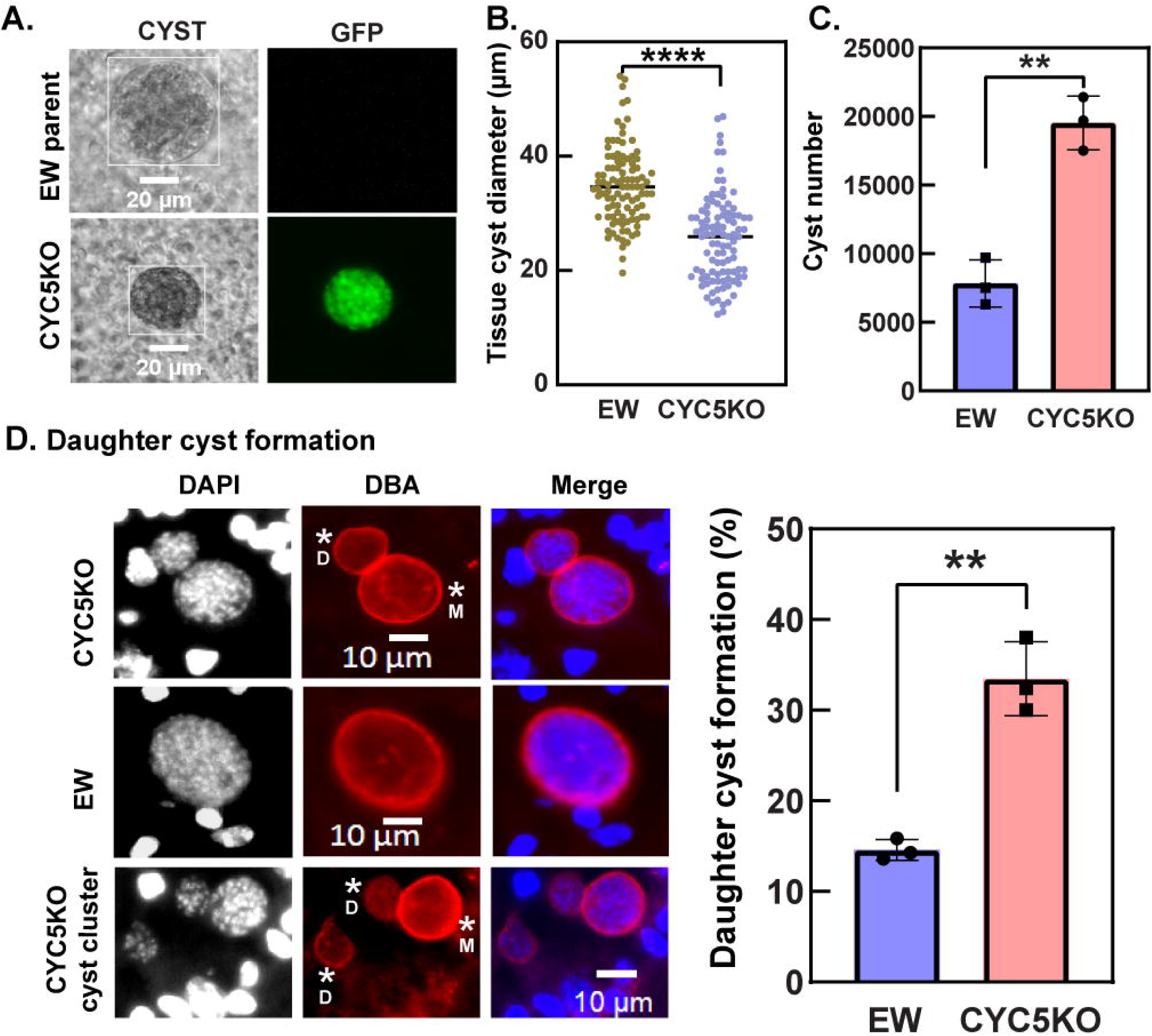
The loss of TgCYC5 increases daughter tissue cysts in vivo. **A**. Comparison of cyst formation in ME49EW::Δ*cyc5* (CYC5KO) and ME49EW (EW) parasites. Only CYC5KO cysts expressed GFP. A 20 µm scale bar is indicated. **B.** Tissue cyst sizes (diameter) were measured using ImageJ software. A total of 250 cysts were cloned in 96-well plates, and images were captured using a microscope prior to measurement. ME49EW cysts formed significantly larger cysts compared to CYC5KO. **C.** CYC5KO parasites produce significantly higher cyst numbers in mouse brain tissue. **D.** Infected brains were cryosectioned and stained with DBA. Large mother cysts are labeled as ‘M’, while small daughter cysts were labeled as ‘D’. Tissue cysts from stained slices were quantified to determine the frequency of daughter cyst formation. Only cysts in close proximity to larger mother cysts were counted as daughter cysts. CYC5KO tissue cysts show significantly increased daughter cyst formation and there were many clusters nearby mother cysts indicating an ongoing daughter cyst production. DBA staining confirmed significantly higher percentage of daughter cysts adjacent to mother cyst, which likely underestimates the number of daughter cysts. DAPI was used to stain host cell and parasite nuclei. Scale bar=10µm.

### Knockout of AP2IX-9 and CYC5 genes inversely alter bradyzoite subtypes in vivo

We have recently developed a new antibody reagent against the surface antigen SRS22A [26], which is expressed in bradyzoites from early to chronic stage tissue cysts (>14 days) in vivo. Expression or lack of SRS22A differentiates two in vivo bradyzoite subtypes that are functionally distinct; SRS22A-bradyzoites primarily initiate the bradyzoite-to-bradyzoite pathways, while SRS22A+ parasites are responsible for bradyzoite conversion to the tachyzoite stage [26]. Here we show that developmental pathways (brady-brady vs brady-tachy) initiated by bradyzoites lacking AP2IX-9 or TgCYC5 were distinctly altered, therefore, we investigated SRS22A expression in ME49EW::Δ*cyc5* versus ME49EW::Δ*ap2IX-9* tissue cysts from chronically infected mice. Thin smears of brain homogenates (30 μl) were fixed, stained with anti-SRS22A antibodies and DBA and the proportion of SRS22A+ cysts calculated (Fig. 7). We expected the fraction of SRS22A-cysts to be affected in ME49EW::Δ*ap2IX-9* bradyzoite infections due to the reduction in bradyzoite-to-bradyzoite replication during ex vivo recrudescence in astrocytes (Fig. 5). As previously seen, ME49EW cysts were composed of SRS22+ and SRS22-expressing cysts with on average 20% of cysts showing no expression of SRS22A (Fig. 7A). In contrast, SRSR22A was detected in all cysts from 28-Day ME49EW::Δ*ap2IX-9* infected mice (Fig. 7A) with a doubling of the cysts that were 100% SRS22A+. Half the cysts in the ME49EW::Δ*ap2IX-9* infections harbored a mixture of SRS22A+ and SRS22A-bradyzoites (Fig. 7A, intracyst areas lacking anti-SRS22A staining have numerous DAPI staining parasite nuclei), however, 100% SRS22A-cysts were completely absent.

**Figure 7.**
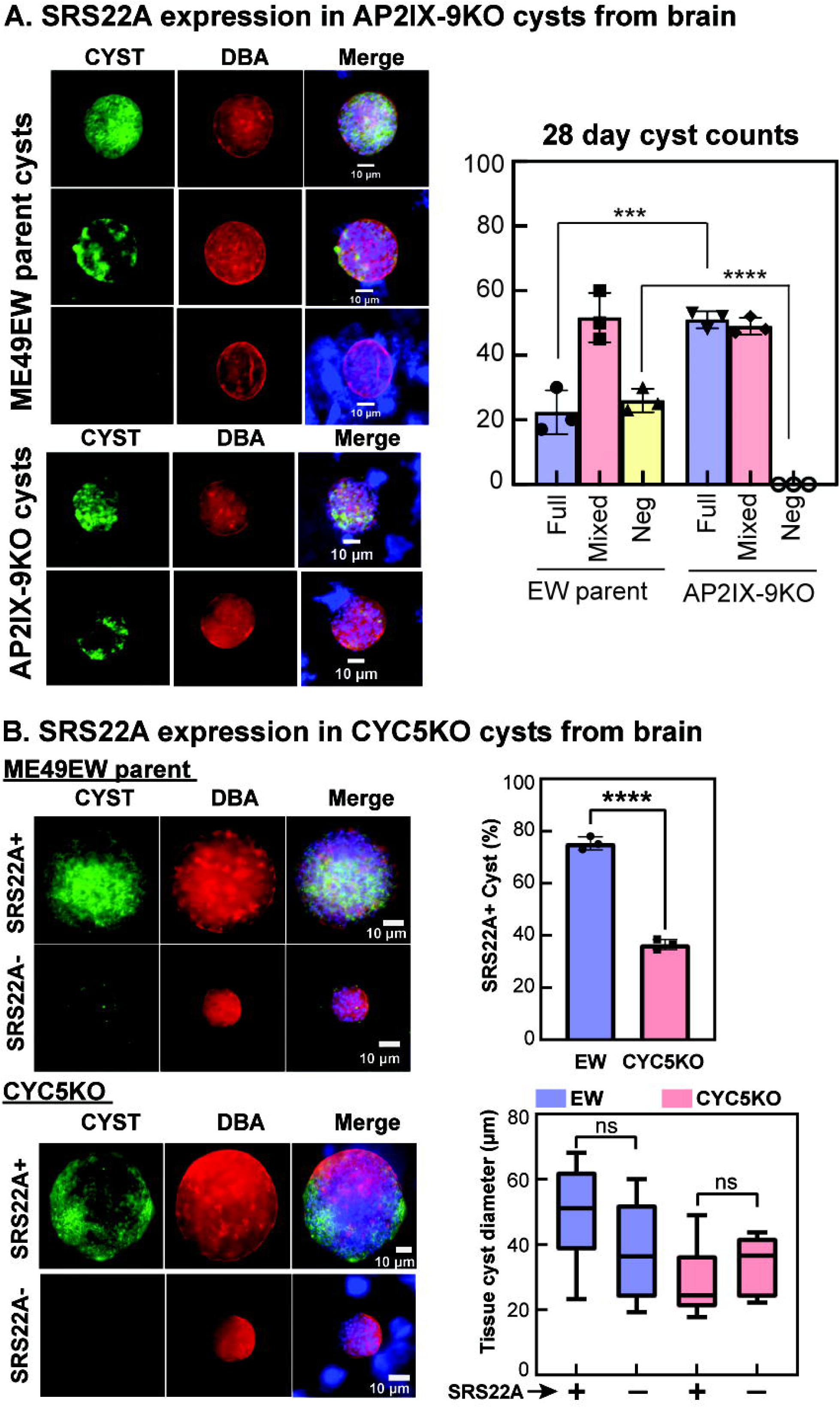
SRS22A bradyzoite subtypes are influenced by genetic deletion of TgCYC5 or ApiAP2IX-9. SRS22A expression in tissue cysts from 28-30 Day infected mouse brain. Infected brain tissue homogenates were fixed and stained for SRS22A expression (anti-SRS22A antiserum) and cyst wall (DBA+). SRS22A+ is shown in green and DBA+ is shown in red. Dapi stained host cell and parasite DNA are included in the merge images. Scale bars are included for representative cyst images. **A.** All tissue cysts from mice infected with ME49EW::Δ*ap2IX-9* (AP2IX-9KO) bradyzoites harbored SRS22A+ bradyzoites (equal 100% SRS22A+ or mixed SRS22A+/SRS22A-cysts). No SRS22A-cysts were detected, while EW parent infections produced >20% SRS22A negative cysts (see graph). Representative images of full, mixed, and negative cysts are shown for EW parent and AP2IX-9KO 28-Day bradyzoite infections. **B.** The majority of the tissue cysts in the EW parent infected mice were were positive for SRS22A (∼75% were full or partially SRS22A+, top graph). By contrast, ME49EW::Δ*cyc5* (CYC5KO) parasites failed to produce the number of SRS22A+ tissue cysts (∼65% of CYC5KO cysts were SRS22A-) as compared to the EW parent Note that the small tissue cyst size of CYC5KO infections were confirmed (see Fig. 6), however, cyst size did not appear to influence SRS22A positivity or lack of SRS22A expression (bottom graph).

The balance of SRS22A expression in ME49EW::Δ*cyc5* cysts was shifted in an opposite manner compared to ME49EW::Δ*ap2IX-9* bradyzoite infections, consistent with the early re-emergence of bradyzoites during ME49EW::Δcyc5 ex vivo recrudescence (Fig. 6). The majority (∼65%) of ME49EW::Δ*cyc5* tissue cysts in 30-Day infected mice were SRS22A-, which was a significant change in bradyzoite subtype balance when compared to the ∼75% SRS22A+ cysts formed in ME49EW parental strain (Fig. 7B). In this experiment, we again confirmed that ME49EW::Δ*cyc5* tissue cysts were considerably smaller than ME49EW parent tissue cysts, although SRS22A expression did not correlate to a specific range of cyst size. The distinct shift of ME49EW::Δ*cyc5* parasites to a majority SRS22A-bradyzoites is likely responsible for the increased daughter cyst formation associated with the TgCYC5 knockout in vivo (Fig. 6D).

## Discussion

*Toxoplasma* is a valuable model organism for studying the basic biology of single cell parasitic eukaryotes. There have been steady advances in genetic methods for *Toxoplasma* since the first transient and stable transfection of tachyzoites ∼30 years ago [27, 28]. An expanded list of positive and negative selectable markers now includes seven drug/gene pairs providing positive and/or negative genetic selection [28–31]. Gene replacement and DNA transformation frequencies have been improved by eliminating nonhomologous DNA repair and targeting specific chromosome breaks in or near target genes [32–34]. Because genetic deletion is unsuitable for essential genes, conditional knockdown methods have been developed that operate transcriptionally or post-translationally [35, 36]. Unfortunately, the genetic “tool kit” is tachyzoite-centric and relies on the repeated passage of lab-adapted strains in HFF host cells. It is common knowledge these conditions adversely affect *Toxoplasma* developmental competency (Fig. 1 and 2) [14]. We have sought to solve this problem so that genetic tools can be applied in an unbiased way to the study of *Toxoplasma* developmental biology. 1) We have established protocols for maintaining competent tissue cysts in mice and produce in vivo bradyzoite yields suitable for genetic experiments. 2) Our transfection protocols take advantage of the fast-growing tachyzoite formed by ex vivo bradyzoite infections of primary astrocytes. Fast-growing tachyzoites transfected at frequencies equivalent or better than Type 1 RH tachyzoites. 3) We experimentally verified the clonality of tissue cysts and developed protocols for isolating single cysts and preserving transgenic cysts.

The new ex vivo bradyzoite-based genetic protocols we developed are primed for further advancement (e.g. adding conditional knockout and expression), although by comparison to tachyzoite protocols these experiments are more challenging and take longer to accomplish. Obtaining transgenics parasites can be accomplished in a few weeks using lab-adapted strains like RH, whereas clonal transgenic tissue cysts will take several months to acquire. However, as we and others have shown *Toxoplasma* transgenic strains produced by repeated passage in cell culture are not able to fully replicate *Toxoplasma* in vivo development [14, 37]. Our recent discovery of in vivo SRS22A differential expression and the distinct developmental functions it signals [26] is likely not the only biology missed when using cell culture-adapted strains. Importantly, the failure of in vitro models is not just a loss of capability, but these models can also lead to wrong answers. Our studies of the transcription factor AP2IX-9 in HFF in vitro models determined that overexpression of AP2IX-9 repressed the alkaline-stress induction of some bradyzoite gene expression, and also inhibited alkaline-stress induced cyst wall formation [20]. From these results, we concluded AP2IX-9 must function to negatively regulate tachyzoite-to-bradyzoite development, although at the time we were unable to verify this mechanism in a competent in vivo model of *Toxoplasma* development. Later studies determined that deletion of the AP2IX-9 gene did not inhibit DBA+-cyst wall formation in vitro, while it modestly enhanced cyst wall formation [19]. Targeted CRISPR screens of DNA binding proteins, also determined that AP2IX-9 was not essential for tissue cyst formation in vitro [19]. It was, therefore, not surprising that the deletion of the AP2IX-9 gene in the developmentally competent ME49EW strain using ex vivo bradyzoite-based methods, maintained cyst burden in the brain (Fig. 4). This left us with the question of why AP2IX-9 is the highest expressed ApiAP2 mRNA in chronic stage tissue cysts [14]. The bradyzoite is the multipotent stage of the *Toxoplasma* life cycle; bradyzoite-to-tachyzoite, bradyzoite-to-bradyzoite, and bradyzoite-to-merozoite are pathways that can be initiated by bradyzoite infections. It is clear some of these pathways are host-dependent whereas others are not [14, 38, 39]. Importantly, while deletion of AP2IX-9 in ME49EW tissue cysts did not block bradyzoite-to-tachyzoite recrudescence, it completely eliminated the pathway of bradyzoite-to-bradyzoite replication that is active in ME49EW parent infections of astrocytes but not fibroblasts [14]. The function of direct bradyzoite replication is not fully understood. This developmental process may be required for long term maintenance of the tissue cysts in animals, or it may prepare the bradyzoite for transmission to different animal hosts, which were functions not evaluated here. The role of AP2IX-9 in bradyzoite-to-merozoite development was also not investigated. Further studies will be needed to fully understand the role of AP2IX-9 in bradyzoite developmental biology and its role in the initiation of the sexual stage pathways in the feline gut. The developmentally competent transgenic cysts lacking AP2IX-9 generated in this study now make these studies possible.

*Toxoplasma* cyclin proteins have not been studied in the context of bradyzoite recrudescence, however, they have been characterized in tachyzoites by various groups using cell culture-adapted *Toxoplasma* strains [25, 40, 41]. This prompted us to investigate the role of the bradyzoite-specific cyclin, TgCYC5, that is highly expressed in tissue cysts within the mouse brain and confirm the ability of our ex vivo bradyzoite methodology to investigate cyst biology. A previous study of TgCYC5 that employed an auxin-inducible conditional knockdown system, reported a significant reduction in tissue cyst formation in rat neuron cultures when TgCYC5 was destabilized [25]. Thus, the in vitro phenotype of TgCYC5 conditional knockout indicated this bradyzoite cyclin was required if not essential for tissue cyst development. To reexamine this role for TgCYC5, we generated a TgCYC5 knockout in the developmentally competent ME49EW strain using the bradyzoite-centric protocols introduced here. Surprisingly, ME49EW::Δ*cyc5* parasites readily formed DBA-positive tissue cysts in mice at levels substantially higher than the ME49EW parent strain (Fig. 6). Consistent with the higher cyst burden in mice, the loss of TgCYC5 in the ex vivo bradyzoite recrudescence model led to a premature shift of the fast-growing tachyzoite to slower growth and the early re-emergence of bradyzoites (Fig. 5). In ME49EW::Δ*cyc5* chronic infections of mouse brain, this “short-circuiting” of bradyzoite-to-tachyzoite recrudescence caused an increased presence of daughter cysts and numerous clusters of small cysts (Fig. 6). Thus, similar to the AP2IX-9 quandary, in vitro versus in vivo developmental models uncovered contradictory roles for TgCYC5 in bradyzoite development.

Our recent discovery of bradyzoite subtypes with distinct functions [26] may now provide an explanation for how TgCYC5 and AP2IX-9 influence *Toxoplasma* development. We have recently established a new developmental paradigm based on the discovery of multiple bradyzoite subtypes and the demonstration that the SRS22A-bradyzoite subtype is primarily responsible for initiating the bradyzoite-to-bradyzoite pathway and the SRS22A+ bradyzoite subtype initiates the bradyzoite-to-tachyzoite pathway [26]. The deletion of AP2IX-9 or TgCYC5 distinctly altered the balance of direct bradyzoite replication versus conversion of bradyzoites to the tachyzoite stage in the ex vivo recrudescence assay (Figs. 4 and 5). This new model (Fig. 8) successfully predicts the balance of SRS22A expression in the ME49EW::Δ*cyc5* and ME49EW::Δ*ap2IX-9* strains (Fig. 7). The normal composition of SRS22A bradyzoite subtypes in ME49EW infections favors SRS22A+ tissue cysts (∼75-80%; Fig. 7), which primarily initiate bradyzoite-to-tachyzoite development [26]. Genetically manipulating AP2IX-9 further shifts the balance towards SRS22A+ cysts with no SRS22A negative cysts detected when the AP2IX-9 gene was deleted indicating a surprising role for AP2IX-9 in preserving the bradyzoite-to-bradyzoite pathway (Fig. 8). By contrast, the loss of TgCYC5 dramatically shifts the developmental balance towards SRS22A-bradyzoites (∼65, Fig. 7B), which favors direct bradyzoite-to-bradyzoite replication [26] and is likely responsible for the increased production of small daughter cysts in mice infected with the ME49EW::Δ*cyc5* transgenic strain. ME49EW::Δ*cyc5* bradyzoites reduced capacity to form SRS22A+ bradyzoites also caused the reduction in the bradyzoite-to-tachyzoite pathway we observed in the ex vivo recrudescent model and abridged the duration of the fast-growing tachyzoite stage (Fig. 5). Intriguingly, the small cyst size and majority SRS22A-bradyzoite profile of ME49EW::Δ*cyc5* parasites at 30-day infection in mice is very similar to Day-14 tissue cysts in ME49EW parent infections [26]. Therefore, it is possible that the loss of TgCYC5 caused ME49EW::Δ*cyc5* parasites to become blocked at an early stage of tissue cyst development or alternatively they are developing much more slowly in mice. Altogether, the ex vivo and in vivo studies of ME49EW::Δ*cyc5* bradyzoites indicates TgCYC5 cyclin has an unexpected role in augmenting the number of SRS22A+ bradyzoites that would enhance conversion to tachyzoites (Fig. 8). More work will be required to fully understand these complex mechanisms active in the *Toxoplasma* intermediate life cycle. Fortunately, the ME49EW::Δ*ap2IX-9* and ME49EW::Δ*cyc5* strains provide new avenues to study how the TgCYC5 cyclin and its kinase partner TgCrk2 [25] and the transcription factor AP2IX-9 operate to control the balance of bradyzoite subtypes during *Toxoplasma* development in the intermediate host life cycle.

**Figure 8.**
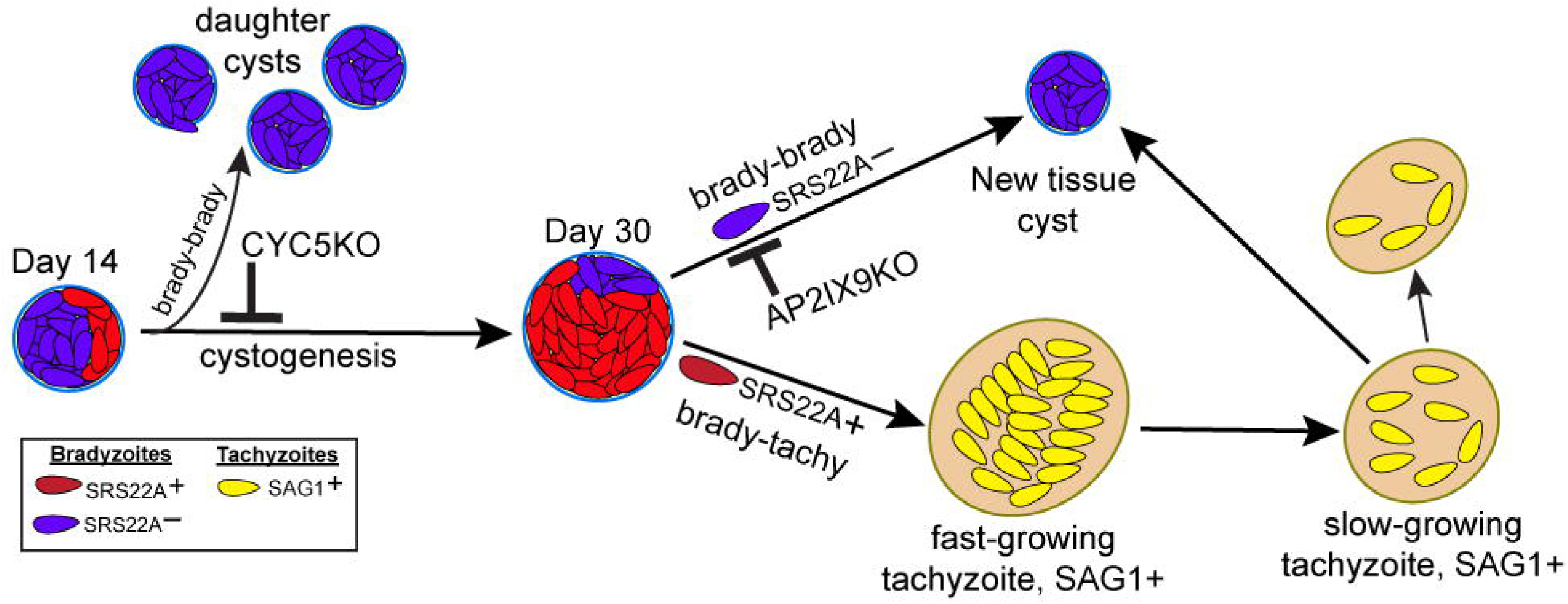
Bradyzoite subtypes have critical roles in *Toxoplasma* development. Cystogenesis of the Type II ME49EW strain shows progressive development of two bradyzoite subtypes in mice with the SRS22A-subtype the majority early and the SRS22A+ subtype more prevalent in the later chronic disease phase. Recent studies demonstrate that SRS22A-bradyzoites primarily initiate direct bradyzoite replication, whereas SRS22A+ bradyzoites are responsible for bradyzoite to tachyzoite conversion [26]. The loss of TgCYC5 cyclin shifts development towards the bradyzoite replication pathway leading to increased daughter cysts, whereas deletion of the transcription factor TgAP2IX-9 shifts the developmental balance further towards the SRS22A+ subtype during the chronic disease phase.

In summary, we have developed ex vivo bradyzoite protocols that were combined with the current *Toxoplasma* genetic “tool kit” to enable for the first time the generation of transgenic tissue cysts in mice. The critical parasites of this new genetic model are the unadapted ME49EW strain from which developmentally competent bradyzoites from in vivo tissue cysts can be purified and via ex vivo bradyzoite infections the unique fast-growing tachyzoite capable of efficient transformation can be produced. We have used these new protocols to revisit the accuracy of AP2IX-9 and TgCYC5 mechanisms proposed by in vitro models of bradyzoite development, while at the same time discovering new mechanisms that were inaccessible in conventional cell culture models of bradyzoite development.

## Materials and Methods

### Parasite culture and host cells

Type II ME49EW developmentally competent strain was used in this study. ME49EW bradyzoites were seeded on monolayers of murine astrocytes from ScienCell (#M1800) as per manufacturer’s protocols or isolated from C57BL/6 mouse pups born in our vivarium facility as previously described [14].

### Animal protocols

Infections were performed in 6 weeks old female CBA/j or Swiss Webster (SWR) mice purchased from Jackson laboratories and Taconic biosciences respectively. ME49EW parent strain and other transgenic strains was sequentially passaged in CBA/j and SWR/j mice to preserve the developmental competency to form tissue cysts at high numbers. Ten cysts/200 µl from SWR/j brain cortex were used to infect CBA/j mice and then 30-60 days post-infection 10 cysts/200 µl from CBA/J brains were used to re-infect SWS/j mouse strains to continue the process. All mice were maintained at UCR vivarium facility strictly following animal protocols from IACUC (Institutional Animal Care and Use Committee). In the current studies, maintenance of strains was maintained by alternating SWR/j and CBA/j mice infections. In addition, SWR/j mice were used for generating the Day-2 fast-tachyzoites needed for transfections. CBA/j mice were used for testing cyst production and for evaluating the biology of knockout strains. We previously showed that ME49EW cyst numbers in CBA/j infected mice increase between 30 and 40 d.p.i. [14]. For production purposes, a 5-10% mortality by 40 days post-infection was factored for ME49EW strain infections of CBA/j mice. Doses higher than 10 cysts can be used, however, increases in cyst yields per mouse were offset by higher CBA/j mortality.

### Bradyzoite recrudescence experiments

ME49EW bradyzoites from mice were harvested and purified and used to infect confluent primary astrocytes grown under 5% oxygen (see Protocol S1 for full details). For immunofluorescent studies, excysted bradyzoites were seeded at 0.5 multiplicity of infection on poly-L-Lysine coated coverslips of primary astrocytes. At various times recrudescing parasites were fixed for immunofluorescence experiments. To avoid host cells becoming a limiting factor, and to allow for continuous tracking of the growth rate, parasites were passed post ex vivo infection after Day 3 and Day 5 (and for additional experiments after Day 7) prior to host cell lysis [16].

### Staining of tissue cysts in the brain lysate

Brain cortex in 3ml 1XPBS were sequentially syringe lysed with 18-, 20-, and 22-gauge hypodermic needles for five times each. 30 µl brain lysate was used to count the number of total cysts in whole brain described elsewhere. Brain lysates were smeared, air dried on the glass slides following fixation in methanol for 5 seconds. Slides were dried and blocked with 5% donkey serum for 30 minutes following treatment with primary antibodies in 5% donkey serum at 4 °C overnight. Next day, slides were washed with 1XPBS and stained with a mixture of DAPI, Dolichos-biflorous (rhodamine) and fluorophore labelled secondary antibodies for 1 hour. Finally, slides were washed with 1XPBS and mounted with mounting media.

### Immunofluorescence experiments

Immunofluorescence assays were conducted as described [14]. Parasites growing on human foreskin fibroblast (HFF) monolayers or murine astrocytes were fixed with freshly prepared 4% paraformaldehyde for 15 minutes at room temperature and washed with phosphate-buffered saline (1xPBS). Fixed cells were then with 100% acetone for 10 minutes following washing with 1xPBS and blocked in 5% donkey serum for 1 hour at room temperature. Primary antibodies diluted in 5% donkey serum were added to the cells, which were incubated overnight at 4°C. After thorough washing with 1X PBS, cells were stained with DAPI (1:1000; Invitrogen) and Alexa Fluor-conjugated secondary antibodies for 1 hour at room temperature. Cells were finally washed with 1XPBS and mounted in ProLong Gold Antifade (Invitrogen) mounting media. The primary antibodies and their respective dilutions used in these experiments were: rabbit α-SRS9, mouse α-SAG1, and rabbit α-SRS22A. Rhodamine conjugated *Dolichos biflorus* lectin stain (1:300; Vector Laboratories) was added directly in the secondary antibody mixture for 1 hour at room temperature to stain cyst wall. Slides were scanned with Leica and Keyence microscopes. The latest software versions for both microscopes were utilized for image analysis.

### HXGPRT and AP2IX-9 knockout with CRISPR-CAS9 system

For disruption of the HXGPRT (TGME49_200320) and AP2IX-9 (TGME49_306620) target genes, we used the multi-guide-RNA (multi-gRNA) CRISPR-Cas9 system as previously published [42]. The CRISPR-Cas9 gRNA plasmid (pSAG1::Cas9-U6::sgUPRT) was provided by David Sibley (Washington University, St. Louis, MO). To generate knockouts in ME49EW Day-2 parasites, three gRNA plasmids were used for the HXGPRT gene, and two gRNA plasmids were used for AP2IX-9 gene. To construct knockout plasmids, a common gene replacement plasmid containing a pyrimethamine resistance gene (TgDHFR^pyr^) was constructed using 3-fragment Gateway protocols (Thermo Fisher Scientific, Waltham, MA). We then PCR amplified a DNA fragment from a plasmid kindly provided by John Boothroyd (Stanford, CA) [43] that encodes a dual expression cassette of tub driven firefly luciferase and gra2 driven GFP. The GFP-Fluc cassette was then inserted into the TgDHFR^pyr^ entry vector using In-Fusion HD cloning (Takara Bio.) that resulted in the triple gene entry plasmid (GFP_Fluc_TgDHFR^pyr^), which was used for all target gene replacements. To construct specific knockout plasmids, the 5’ and 3’ untranslated regions (UTRs) of the HXGPRT or AP2IX-9 genes were PCR amplified from Type II parasite genomic DNA (see primer designs below) and the resulting fragments cloned into Gateway entry plasmids. The 5’ UTR of each target gene was cloned into the pDONR_P4-P1r vector, and the 3’ UTR was cloned into the pDONR_P2r-P3 vector using the BP recombination reaction. After isolation of the pDONR_P4-P1r_5’UTR and pDONR_P2r-P3_3’UTR plasmids for each target gene, they were combined with the GFP_FLuc_TgDHFR^pyr^ plasmid in an LR recombination reaction to generate a final knockout plasmid for each target gene. The final knockout plasmids were linearized before introduction into ME49EW parasites by nucleofection.

Table of primers used to construct guide RNA plasmids and left and right arm entry vectors for genes TgHXGPRT and AP2IX-9.

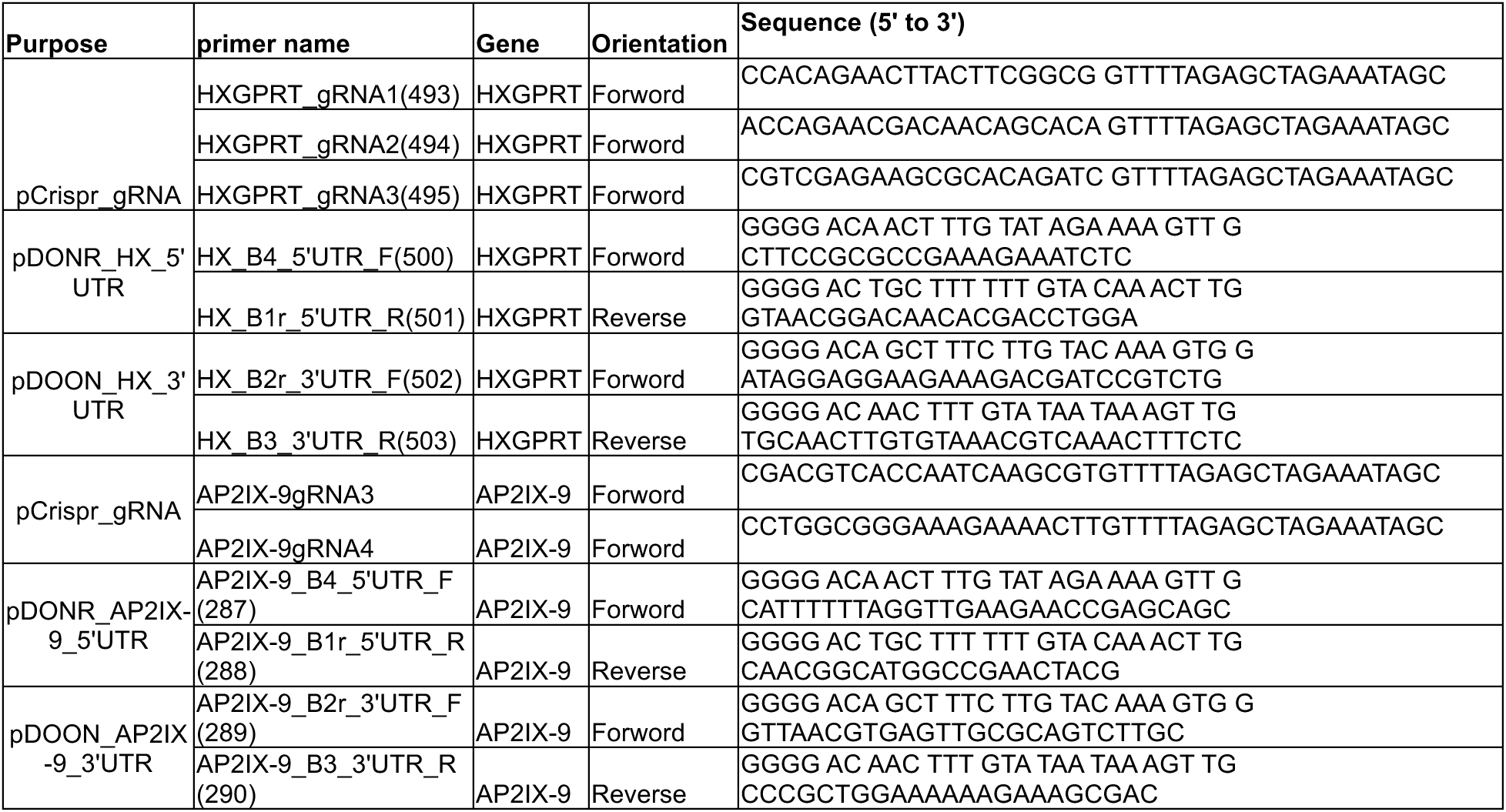

### Transfection and cloning methods (see Protocol S2 for full details)

To preserve cystogenic potential, ME49EW parent tissue cysts and transgenic lines were maintained cyclically in CBA/J and SWR mouse strains. Bradyzoite cysts purified from CBA/J mouse brains were treated in-vitro with pepsin-HCl to mimic natural excystation in the acidic environment of the stomach, allowing the isolation of acid-resistant bradyzoites. The excysted bradyzoites were then cultured on primary murine astrocytes under 5% O₂ (low) to recreate a brain-like environment appropriate to bradyzoite growth and recrudescence. Notably, these conditions were successfully achieved by utilizing gas chambers commonly used for malaria merozoite propagation providing a robust and reliable platform for short-term cultivation of recrudescence populations. Parasites from Day-2 astrocyte cultures (post-ex vivo bradyzoite infection) were nucleofected with a combination of CRISPR-guide plasmids and GFP-equipped drug-resistant cassettes, and then selected under pyrimethamine (0.5μM) for 72 h. Drug-selected parasites were FACS-sorted for GFP+ parasites and injected i.p. into CBA/j mice within 7 days post-bradyzoite excystation to avoid loss of developmental stages. GFP+ tissue cysts were isolated 30-40 days post-infection and cloned by limiting dilution.

### Detailed protocols in supplement

Protocol S1: Cyst and bradyzoite purification

Protocol S2: Producing transgenic cysts

Protocol S3: Cloning transgenic tissue cysts.

Protocol S4: Freezing cysts in brain homogenates.

## Supporting information

Protocol S1

Protocol S2

Protocol S3

Protocol S4

## Acknowledgements

*Toxoplasma gondii* gene information was accessed on http://ToxoDB.org. We thank Dr. John Boothroyd for antibodies and plasmids used in our study and Dr. David Sibley for the *Toxoplasma* adapted CAS9 plasmid. We would like to acknowledge Loic Ciampossin and Dr. Karine LeRoch for their help in validating the genetic knockout strains. This study was supported by NIH NIAID R01 grants AI158417 and AI124682 to M.W.W. and E.H.W., AI122760 to M.W.W., and DA048815 to E.H.W.

## References

1. Brasil, T.R., et al., Host-Toxoplasma gondii Coadaptation Leads to Fine Tuning of the Immune Response. Front Immunol, 2017. 8: p. 1080.

2. Sullivan, W.J., Jr. and V. Jeffers, Mechanisms of Toxoplasma gondii persistence and latency. FEMS Microbiol Rev, 2012. 36(3): p. 717–33.

3. Basavaraju, A., Toxoplasmosis in HIV infection: An overview. Trop Parasitol, 2016. 6(2): p. 129–135.

4. Guex-Crosier, Y., Update on the treatment of ocular toxoplasmosis. Int J Med Sci, 2009. 6(3): p. 140–2.

5. Martynowicz, J., J.S. Doggett, and W.J. Sullivan, Jr., Efficacy of Guanabenz Combination Therapy against Chronic Toxoplasmosis across Multiple Mouse Strains. Antimicrob Agents Chemother, 2020. 64(9).

6. Alday, P.H. and J.S. Doggett, Drugs in development for toxoplasmosis: advances, challenges, and current status. Drug Des Devel Ther, 2017. 11: p. 273–293.

7. Dunay, I.R., et al., Treatment of Toxoplasmosis: Historical Perspective, Animal Models, and Current Clinical Practice. Clin Microbiol Rev, 2018. 31(4).

8. Yan, J., et al., Meta-analysis of prevention and treatment of toxoplasmic encephalitis in HIV-infected patients. Acta Trop, 2013. 127(3): p. 236–44.

9. Aspinall, T.V., et al., The molecular basis of sulfonamide resistance in Toxoplasma gondii and implications for the clinical management of toxoplasmosis. J Infect Dis, 2002. 185(11): p. 1637–43.

10. Montazeri, M., et al., Drug Resistance in Toxoplasma gondii. Front Microbiol, 2018. 9: p. 2587.

11. Barrett, M.P., et al., Protozoan persister-like cells and drug treatment failure. Nat Rev Microbiol, 2019. 17(10): p. 607–620.

12. Waldman, B.S., et al., Identification of a Master Regulator of Differentiation in Toxoplasma. Cell, 2020. 180(2): p. 359–372 e16.

13. Sokol-Borrelli, S.L., et al., A transcriptional network required for bradyzoite development in Toxoplasma gondii is dispensable for recrudescent disease. Nat Commun, 2023. 14(1): p. 6078.

14. Vizcarra, E.A., et al., An ex vivo model of Toxoplasma recrudescence reveals developmental plasticity of the bradyzoite stage. mBio, 2023. 14(5): p. e0183623.

15. Boillat, M., et al., Neuroinflammation-Associated Aspecific Manipulation of Mouse Predator Fear by Toxoplasma gondii. Cell Rep, 2020. 30(2): p. 320–334 e6.

16. Ndubuizu, O. and J.C. LaManna, Brain tissue oxygen concentration measurements. Antioxid Redox Signal, 2007. 9(8): p. 1207–19.

17. Lemgruber, L., et al., The organization of the wall filaments and characterization of the matrix structures of Toxoplasma gondii cyst form. Cell Microbiol, 2011. 13(12): p. 1920–32.

18. Altschul, S.F., et al., The construction and use of log-odds substitution scores for multiple sequence alignment. PLoS Comput Biol, 2010. 6(7): p. e1000852.

19. Hong, D.P., J.B. Radke, and M.W. White, Opposing Transcriptional Mechanisms Regulate Toxoplasma Development. mSphere, 2017. 2(1).

20. Radke, J.B., et al., ApiAP2 transcription factor restricts development of the Toxoplasma tissue cyst. Proc Natl Acad Sci U S A, 2013. 110(17): p. 6871–6.

21. Radke, J.B., et al., Transcriptional repression by ApiAP2 factors is central to chronic toxoplasmosis. PLoS Pathog, 2018. 14(5): p. e1007035.

22. Wang, J.L., et al., The transcription factor AP2XI-2 is a key negative regulator of Toxoplasma gondii merogony. Nat Commun, 2024. 15(1): p. 793.

23. Antunes, A.V., et al., In vitro production of cat-restricted Toxoplasma pre-sexual stages. Nature, 2024. 625(7994): p. 366–376.

24. Alvarez, C.A. and E.S. Suvorova, Checkpoints of apicomplexan cell division identified in Toxoplasma gondii. PLoS Pathog, 2017. 13(7): p. e1006483.

25. Naumov, A.V., et al., Restriction Checkpoint Controls Bradyzoite Development in Toxoplasma gondii. Microbiol Spectr, 2022. 10(3): p. e0070222.

26. 26. Ulu, A., et al., Bradyzoite subtypes rule the crossroads of Toxoplasma development. BioRxiv, 2025.

27. Kim, K., D. Soldati, and J.C. Boothroyd, Gene replacement in Toxoplasma gondii with chloramphenicol acetyltransferase as selectable marker. Science, 1993. 262(5135): p. 911–4.

28. Soldati, D. and J.C. Boothroyd, Transient transfection and expression in the obligate intracellular parasite Toxoplasma gondii. Science, 1993. 260(5106): p. 349–52.

29. Donald, R.G., et al., Insertional tagging, cloning, and expression of the Toxoplasma gondii hypoxanthine-xanthine-guanine phosphoribosyltransferase gene. Use as a selectable marker for stable transformation. J Biol Chem, 1996. 271(24): p. 14010–9.

30. Messina, M., et al., Stable DNA transformation of Toxoplasma gondii using phleomycin selection. Gene, 1995. 165(2): p. 213–7.

31. Behnke, M.S., A. Khan, and L.D. Sibley, Genetic mapping reveals that sinefungin resistance in Toxoplasma gondii is controlled by a putative amino acid transporter locus that can be used as a negative selectable marker. Eukaryot Cell, 2015. 14(2): p. 140–8.

32. Huynh, M.H. and V.B. Carruthers, Tagging of endogenous genes in a Toxoplasma gondii strain lacking Ku80. Eukaryot Cell, 2009. 8(4): p. 530–9.

33. Fox, B.A., et al., Efficient gene replacements in Toxoplasma gondii strains deficient for nonhomologous end joining. Eukaryot Cell, 2009. 8(4): p. 520–9.

34. Winiger, R.R. and A.B. Hehl, A streamlined CRISPR/Cas9 approach for fast genome editing in Toxoplasma gondii and Besnoitia besnoiti. J Biol Methods, 2020. 7(4): p. e140.

35. Sharifpour, M.F., et al., A GRA2 minimal promoter improves the efficiency of TATi / Tet-Off conditional regulation of gene expression in Toxoplasma gondii. Mol Biochem Parasitol, 2021. 244: p. 111384.

36. Brown, K.M., S. Long, and L.D. Sibley, Plasma Membrane Association by N-Acylation Governs PKG Function in Toxoplasma gondii. mBio, 2017. 8(3).

37. Watson, G.F. and P.H. Davis, Systematic review and meta-analysis of variation in Toxoplasma gondii cyst burden in the murine model. Exp Parasitol, 2019. 196: p. 55–62.

38. Dubey, J.P., Advances in the life cycle of Toxoplasma gondii. Int J Parasitol, 1998. 28(7): p. 1019–24.

39. Martorelli Di Genova, B., et al., Intestinal delta-6-desaturase activity determines host range for Toxoplasma sexual reproduction. PLoS Biol, 2019. 17(8): p. e3000364.

40. Hawkins, L.M., et al., The Crk4-Cyc4 complex regulates G(2)/M transition in Toxoplasma gondii. EMBO J, 2024. 43(11): p. 2094–2126.

41. Hawkins, L.M., et al., Novel CRK-Cyclin Complex Controls Spindle Assembly Checkpoint in Toxoplasma Endodyogeny. mBio, 2021. 13(1): p. e0356121.

42. Shen, B., et al., Efficient gene disruption in diverse strains of Toxoplasma gondii using CRISPR/CAS9. mBio, 2014. 5(3): p. e01114–14.

43. Kim, S.K., A. Karasov, and J.C. Boothroyd, Bradyzoite-specific surface antigen SRS9 plays a role in maintaining Toxoplasma gondii persistence in the brain and in host control of parasite replication in the intestine. Infect Immun, 2007. 75(4): p. 1626–34.

