## Supplemental Figures & Legends for "Mechanisms regulating reactivation pathways of *Toxoplasma gondii* as revealed by bradyzoite transgenesis"

A.

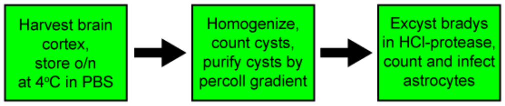


| Me49EW parasite yields | | | | |
|---|---|---|---|---|
| Harvest Date | Days PI | # mice | Bradyzoite yield ($10^6$) | Bradys/ mouse ($10^6$) |
| 7/8/19 | 40 | 20 | 22.0 | 1.1 |
| 1/7/20 | 41 | 22 | 31.3 | 1.4 |
| 7/7/20 | 41 | 20 | 11.9 | 0.6 |
| 8/18/20 | 48 | 20 | 29.6 | 1.5 |
| 12/16/20 | 47 | 20 | 15.7 | 0.8 |
| 3/4/21 | 42 | 20 | 25.7 | 1.3 |
| 11/15/21 | 41 | 15 | 48.6 | 3.2 |
| 8/12/21 | 44 | 25 | 36.1 | 1.4 |
| 8/26/21 | 43 | 11 | 15.5 | 1.4 |
| 9/9/21 | 44 | 15 | 24.2 | 1.6 |
| 11/1/21 | 42 | 22 | 27.6 | 1.3 |
| | | 210 total | 288.2 total | 1.4 average |

B.

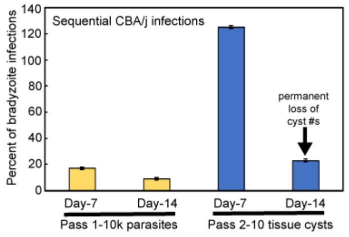


C.

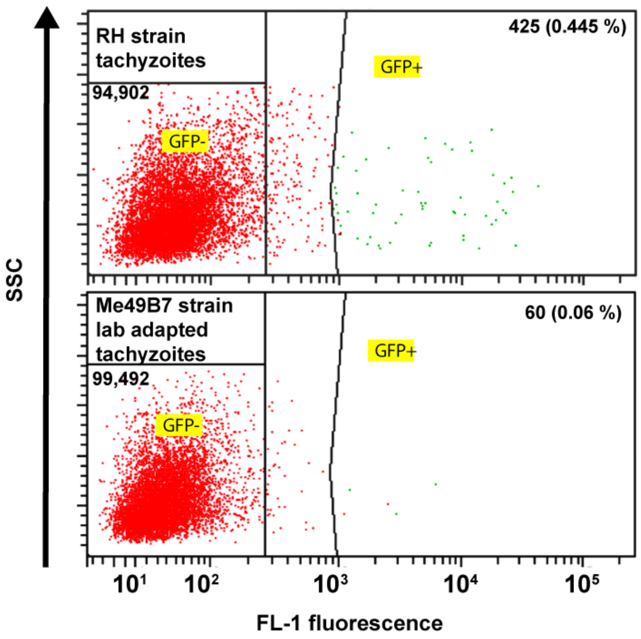


### Figure S1

**A.** Key steps in purifying ME49EW tissue cysts and excysting bradyzoites are indicated. An average of 1.4 million purified bradyzoites are obtained from each CBA/j mouse at 40 d.p.i. (10 cysts i.p.), which is consistent year over year. Detailed procedures for purifying cysts and bradyzoites are included in Protocol S1. **B.** Producing ME49EW tissue cysts and bradyzoites is robust and reproducible. Key steps in purifying ME49EW tissue cysts and excysting bradyzoites are indicated. An average of 1.4 million purified bradyzoites are obtained from each CBA/j mouse at 40 d.p.i. (10 cysts i.p.), which is consistent year over year. Detailed procedures for purifying cysts and bradyzoites are included in Protocol S1. **C.** Transformation efficiency of RH and ME49B7 strain. The TgHXGPRT gene was targeted for CRISPR-assisted knockout in two laboratory strains (RH and ME49B7). Following nucleofection and 96 h post-pyrimethamine (0.5μM) selection in astrocytes, parasites were purified and 100,000 live events/strain analyzed by flow cytometry. Gates defining GFP- versus GFP+ parasites are shown for each strain transfected.

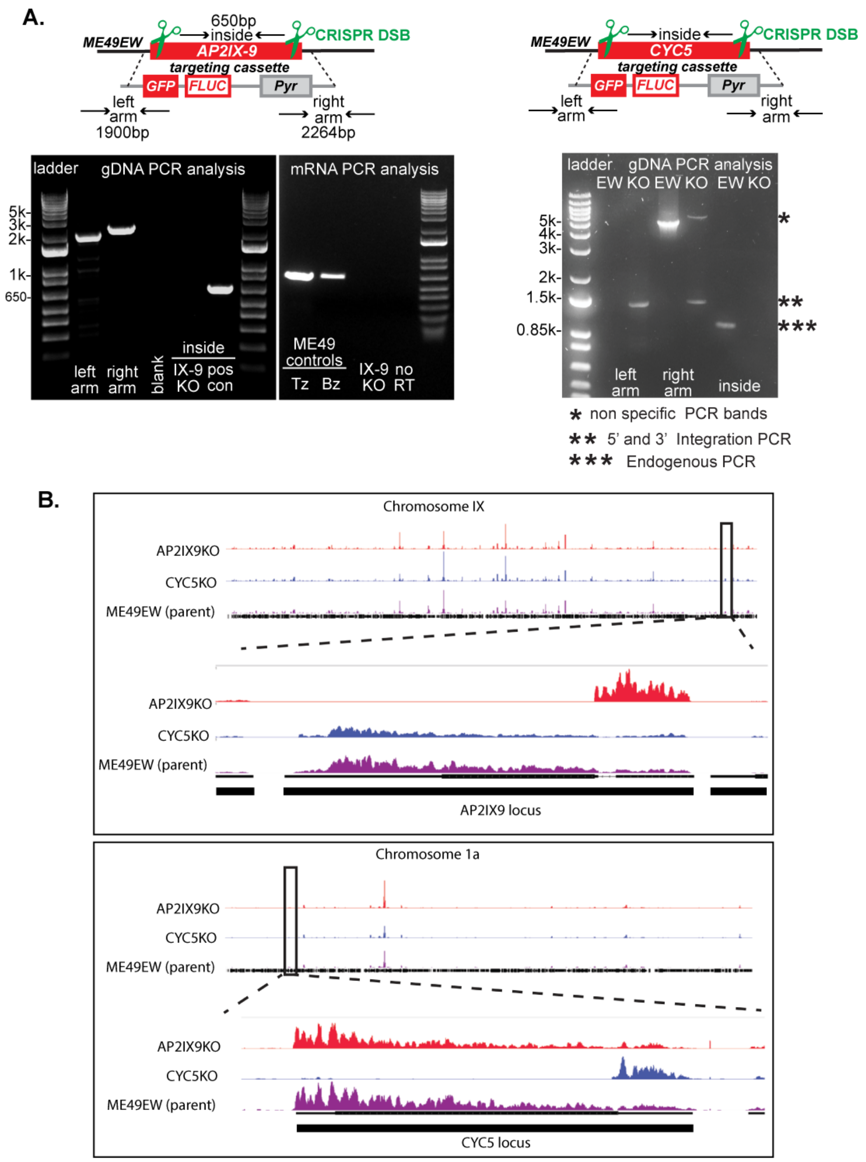


**Figure S2**

**A.** Knockout (KO) of the AP2IX-9 and CYC5 genes in ME49EW tissue cysts were confirmed in a GFP+-tissue cyst cloned by single cyst mouse infection and then expanded in CBA/j mice. The presence of the correct left and right arm DNA fragments in knockout parasite gDNA verified the correct double cross-over had occurred in the clones. The failure to amplify a AP2IX-9 650bp internal gDNA coding fragment as well as the absence of AP2IX-9 mRNA (right gel) was further evidence of the AP2IX-9 gene knockout. RNA purified from ME49EW tachyzoites and cysts provided positive controls for AP2IX-9 mRNA expression. The no reverse transcriptase control (no RT) was included to verify the AP2IX9-KO RNA sample was free of contaminating gDNA. Similarly, CYC5KO failed to amplify a CYC5 1000bp internal gDNA and successfully amplified left and right arms. **B.** Genetic mapping of total RNA-seq reads confirm the loss of the TgAP2IX-9 gene locus on chromosome IX in the ME49EW::Δ*ap2IX-9* strain, which is present in the EW parent strain and the ME49EW::Δ*tgCYC5* strain. This analysis also verified the absence of any AP2IX-9 gene duplication in the knockout strain. In a similar analysis, deletion of the TgCYC5 locus in the ME49EW::Δ*tgCYC5* strain was confirmed on chromosome 1a while eliminating any possible duplication of TgCYC5 gene.
