## Supplementary material for "Mechanisms regulating reactivation pathways of *Toxoplasma gondii* as revealed by bradyzoite transgenesis": Protocol S1

**Protocol S1: Purifying cysts and bradyzoites from mouse brain cortex.**

Preparing brain homogenates

1. At 30-50 days remove the brain cortex using the following procedure:

Remove the head and peel back the skin and membranes from the skullcap exposing the eye sockets. Using the eye sockets hold the head firmly with tooth tissue forceps and remove all soft tissue from what is left of the neck tissue. With sharp scissors, cut along each side of the skull cap (best if someone shows you this step). Carefully pry up the skull cap and remove (you often need to gently reach under the skull cap and push the brain down with your scissors). Slip a flat tool (a scalpel holder without blade works well) under the brain to break the brain stem connection. You should feel a crunch of the bones. While the brain is in place scrape the white lipid and cerebellum away from the cortex. In our studies, the number of cysts in the cerebellum is <10%, which is not worth dealing with the white lipid matter that can clog the needles during homogenization. Drop the cortex into 3 ml of PBS in a 15 ml conical tube on ice (keep brains cold to prevent activating the dormant bradyzoites).

2. Store the brain tissue in 1xPBS overnight at 4°C, which has no deleterious effect on the tissue cysts and greatly improves the homogenization of brain tissue. Homogenizing fresh brain tissue can be done, however, there are a number of drawbacks. The tissue does not homogenize as well and as a result the cysts become trapped in sheets of tissue. This causes problems in the percoll gradient step below and underestimates the number of cysts when counting a 30µl spread by 50% or more.

3. After overnight at 4°C, needle pass the brain tissue with a 18g needle and 3 ml syringe several times. Do the same process sequentially with 20 and 22 g needles until homogenous. Three to four passes per needle gauge is usually enough with post-harvest brain tissue. **Be careful not to create too many bubbles. [start warming up percoll to room temperature before starting homogenization]*

4. Mix the homogenate well and pipette 30 µl onto a microscope slide, and coverslip. Examine using a 5-10x objective under the light microscope to make sure the mouse is infected. To estimate the total number of cysts scan the complete slide: a 5x objective requires 5-6 passes (10 cysts in 30 µl is ~1,000 cysts per cortex). As an example, the field of brain homogenate below contained two tissue cysts.

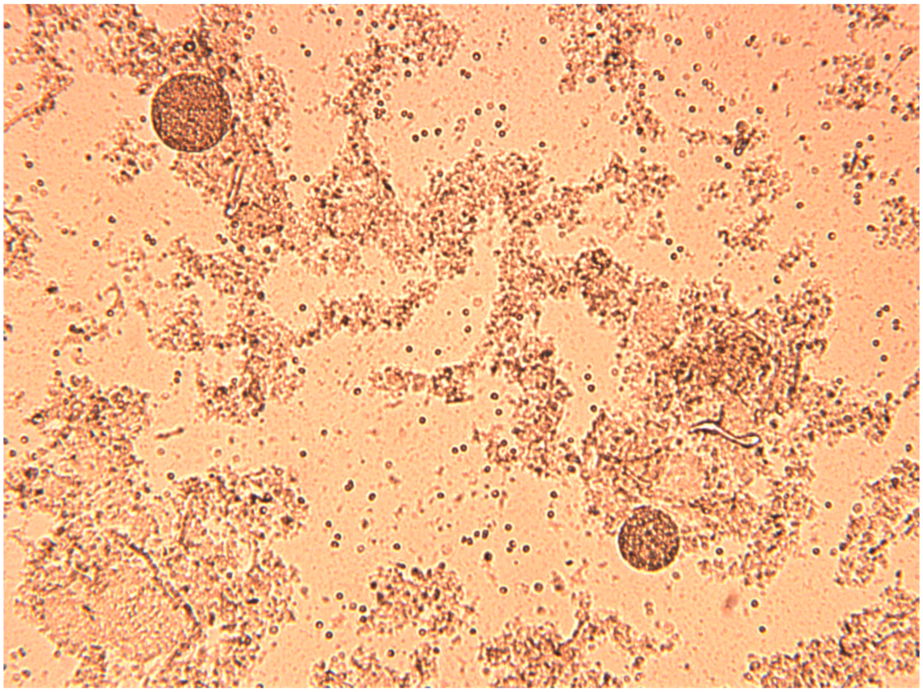

Cyst Purification

5. Add 8 ml of 1x PBS and spin for 15 minutes, 500xg, room temperature. The brain homogenate will pellet (~0.5 ml pellet).

6. During the above spin, prepare 2.5ml room temperature percoll-90 (90% percoll, 10% 1xPBS) for each whole brain sample. **Percoll must be at room temperature.** After spin, pour or aspirate off PBS supernatant and then resuspend the pellet well in 2.5ml percoll-90 by pipetting up and down. Top the tub up to 7.5ml with **room-temp** 1xPBS and invert 3-5 times to mix. (22.5% v/v final in ~0.6x PBS). The lower PBS in the gradient eliminates some of the RBC contamination without harm to the bradyzoites.

7. Spin **room temperature**, **no brakes**, for 15 minutes at 1,000xg. Cysts and RBCs will pellet at the bottom of the tube. **If brakes are on, the gradient will be destroyed. It is imperative the brakes are OFF.*

8. After centrifugation of the percoll gradient, a thick lipid layer should be at the top. RBCs and tissue cysts will be pelleted and there may be a small debris layer above the pellet. Remove the gradient by carefully aspirating all material above the pellet (careful not to suck up the pellet). Remove the thick lipid layer carefully first, then the remaining supernatant. [*Note: you can leave ~100 µl if you are combining pellets*].

9. If there are multiple gradients, combine the pellets at this step (up to 10 gradient pellets can be combined). Add 80 µl of 1x PBS to the first tube, mix and transfer to the next tube (use a P1000 as the volume will increase beyond 200 µl), mix and transfer to the next tube and repeat until you are at the final tube. Wash the tubes with a fresh 80 µl of 1x PBS following this same sequential process. [*Note: at this point the cysts are concentrated, if you take the cyst material out of the hood e.g. need to count, wear a mask*]. Once the pellet material is combined bring up to 10 ml with PBS and spin down. Centrifuge at 30xg for 10 min with a low brake setting. Carefully pour off or aspirate the 1xPBS and resuspend the pellet in 0.2ml of 1xPBS. Check cyst number and contamination on a hemocytometer. Any minimal RBC contamination will be eliminated in the next pepsin step.

Bradyzoite excystation

10. To obtain bradyzoites, add 0.8 ml of pre-warmed pepsin-HCl solution (final 1ml volume), vortex gently and incubate in 37°C water bath for no more than 2 minutes. [*Pepsin-HCl: 50mg pepsin (1:10000 activity), 50 mg NaCl, 70 µl (10 N) to 5 ml pure water. Pepsin final 10mg/ml, NaCl 10mg/ml*]

11. Add warm media (10% FBS) to 12 ml and spin at 2000xg for 10 min. Remove supernatant by aspiration, careful don't go too close to the pellet. Resuspend the pellet in 0.1-0.2 ml media, count bradyzoites and infect host cell cultures.

*Cysts production and purification protocol notes:*

(i). Cyst numbers vary by mouse strain: Sensitivity to the ME49EW strain is highest in CBA/j mice, medium in C57B6, and lowest in BALB/c or Swiss Webster mice. Cyst yields are inversely correlated e.g. CBA/j higher cysts vs BALB/c lower cysts. In CBA/j mice, EW cyst numbers increase progressively with the number increasing beyond 30 days (e.g. 10 cysts infection, ~1,500 cysts/brain at 30 days, ~2,500 cysts at 45 days). In addition, cyst size also increases with time with a corresponding increase in bradyzoite number. Medium cysts have ~500 bradyzoites and large cysts have ~1,500 bradyzoites.

(ii). The ME49EW strain can be maintained for longer than two months in Swiss Webster mice (10 cysts i.p. infections). Brain homogenates from a Swiss Webster mouse diluted to 50 cysts/ml with PBS is used to infect either new CBA/j or Swiss Webster mice based on a 0.2ml i.p. inoculation (10 cysts total). Cysts of the EW strain can be more virulent than tachyzoites of the EW strain. At the 10 cyst level, 5 of 50 CBA/j mice will succumb to the infection over a 40 day period.

(iii). The cyst procedure can be scaled up, however, we have evaluated combining brain homogenates and increasing the size of the percoll gradient (e.g. 2 mice/7.5 ml gradient or 6 mice/45 ml gradient) and in each test, cyst numbers were reduced and contamination increased. The single brain cortex per 7.5ml gradient in a 15 ml conical tube gives the highest yields and lowest contamination. Over several years we have obtained an average of 1.4 million bradyzoites per CBA brain cortex with the ME49EW strain using the above protocol.

(iv). Tips for handling large numbers of mice (e.g. 16 mice): we isolate cysts from each brain in separate percoll gradients After aspiration of the percoll spin, we add 80 µl of 1xPBS to the first of the 10 tubes and resuspend the cysts progressively ending with the 10th tube. Follow with a wash of the tubes progressively with another 80 µl 1xPBS. Total volume ~0.5ml due to the residual 1xPBS left after aspiration. Then each tube is brought up to 10 ml to remove most of the RBCs with a low speed spin. Then the pellet is resuspended in 0.2 ml 1xPBS then 0.8 ml of the pepsin excystation solution is added. Caution: excystation is very fast and 2 min in the pepsin/HCl/NaCl solution in the protocol above is more than adequate for full excystation.

(v). CAUTION, cysts settle very quickly out of suspensions. In the time it takes you to set your pipet down after pipetting that 30 µl sample onto your slide, grab a coverslip and make the wet mount the cysts have already settled to the bottom. The settling effect explains why most of the cysts are located in one half of the slide. Because they are very dense (500-1,500 bradys/cyst) you must constantly mix when you are loading syringes for mouse inoculations or sampling for counting otherwise your infections and counts will vary widely.

(vi). Do not use the ACK lysis buffer to remove RBC contamination, this buffer harms the cysts. Use the low 30xg spin to reduce RBCs in the protocol above. Multiple low speed spins will additively reduce RBC contamination; however, cyst yields will suffer also due to the high dilution effect.
