## Supplementary material for "Mechanisms regulating reactivation pathways of *Toxoplasma gondii* as revealed by bradyzoite transgenesis": Protocol S2

### Protocol S2: Producing ME49EW transgenic tissue cysts.

**MATERIALS NEEDED:**

1. SWR/j brains (harvested 30–40 days PI)
   **NOTE:** 6–8 brains should be sufficient to infect each T75 of astrocytes. Seed each T75 of astrocytes with 5–8 million bradyzoites (BZs). At this MOI, each T75 should have about 50-70 million Day-2 parasites after 48 hours, allowing for 5-6 transfections at 10 million Day-2 parasites per nucleofection.
2. Plasmid insert plus CAS9/gRNA plasmid(s) of interest (5µg EACH); EtOH precipitated (stored @ -20°)
3. Astrocytes–T75 flasks (two for Day 2 parasite production and two T75 flasks for each post-transfection)

Lonza Reagent: P3 Primary Cell Solution (Catalog # V4XP-3012)–100µl/nucleofection

**DAY 2 PARASITE PRODUCTION**

1. Process SWR brains for cysts and excyst bradyzoites following Protocol S1.
2. Count/infect up to 8 million BZs per T75 of astrocytes (change astrocyte media prior to BZ infections)
3. Incubate flasks in hypoxic chamber (5% oxygen) at 37°C for 48 hours

**TRANSFECTION DAY**

**PLASMID & gRNA PREP**

1. Centrifuge pDEST_DHFR plasmid and CAS9/gRNA plasmid(s) at 14.8rpm (max speed) for 30 min at 4°C.
2. Remove supernatant
   **NOTE:** Pour off the bulk of the EtOH solution, then quick spin tubes. Using a P20, carefully remove any residual supernatant, prior to adding the Lonza P3 solution. Let air dry on bench.
3. Vortex pDEST_DHFR plasmid and corresponding CAS9_gRNA plasmid(s) for each transfection in a total of 100µl Lonza P3 Primary Cell Solution. Keep at room temperature after resuspending.

**DAY 2 PARASITE PREPARATION**

1. Scrape, needle pass, filter Day 2 parasites.
2. Determine number of parasites. Aliquot volume containing 10 million parasites and centrifuge at room temperature (1.8xg for 15 minutes).
3. Aspirate supernatant carefully to not disturb the parasite pellet.
4. Resuspend pellet in the 100µl all plasmids solution
5. Transfer entire reaction to a 2mm cuvette
   **NOTE:** Rxn volume should not exceed 125µl (excess volume can come from residual media over the pellet or EtOH from the plasmid/gRNA tubes) – Nucleofector will result in an error message if this reaction volume exceeds the max volume limit.

**TRANSFECTION**

1. Follow Lonza instructions for nucleofection (Nucleofector 4D, X unit). For *Toxoplasma* parasites use program FI-115 (U33 on older nucleofectors).
2. Repeat if performing multiple transfections.

**ONCE TRANSFECTED…**

1. Change media in astrocyte flasks.
2. Add entire transfection reaction to a single T75 astrocyte flask.
3. Incubate flasks in an hypoxic chamber at 37°C without drug selection to let parasites recover.
   Add fresh ScienCell astrocyte media plus 0.5 µM pyrimethamine to flasks 24 h after the recovery period.
4. FACS sort for GFP+ parasites after 4 days of pyrimethamine selection (Day 7 from the original bradyzoite infection).
   **NOTE:** GFP+ parasites are collected between the $10^3$ – $10^5$ gate; see FACS sort example of transgenic ME49 clone positive for GFP expression next page and see also the FACS sort of GFP+ parasites produced by nucleofection of Day-2 ME49EW parasites prior to mouse infection in Figure 4.
5. For each transfection infect at least five CBA/j mice using 5,000 GFP+ parasites/mouse i.p. Adjust as necessary based on the GFP+ sorting numbers.

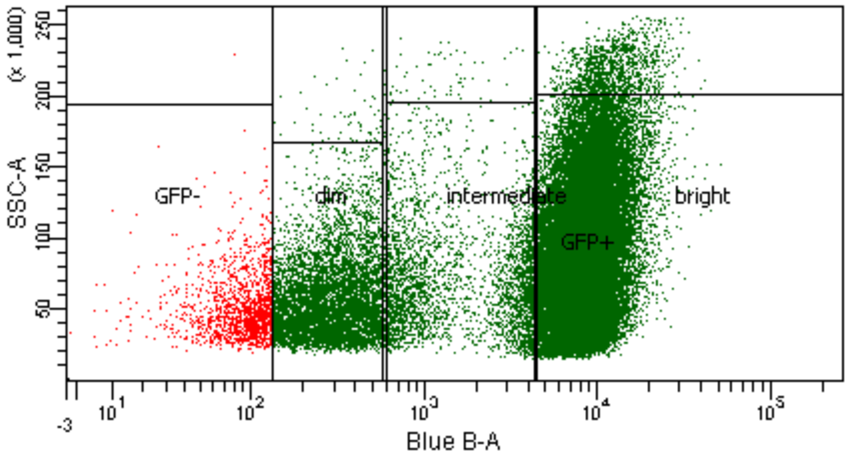


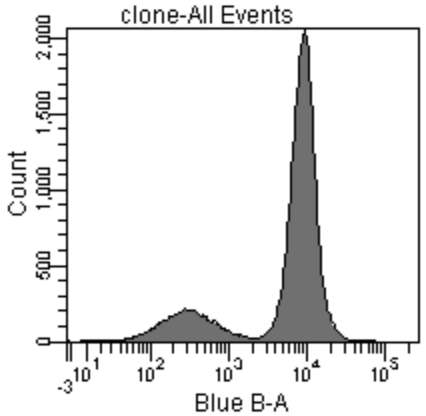


Specimen Name: initial
Tube Name: clone

| Population | %Parent | Blue B-A Median |
|---|---|---|
| P3 | #### | 7,900 |
| GFP+ | 97.2 | 8,021 |
| dim | 12.4 | 284 |
| intermediate | 6.4 | 1,380 |
| bright | 80.5 | 8,844 |
| GFP- | 3.0 | 97 |

Tube: clone

| Population | #Events | %Parent | %Total |
|---|---|---|---|
| All Events | 52,275 | #### | 100.0 |
| P1 | 52,166 | 99.8 | 99.8 |
| P2 | 51,296 | 98.3 | 98.1 |
| P3 | 50,077 | 97.6 | 95.8 |
| GFP+ | 48,660 | 97.2 | 93.1 |
| dim | 6,035 | 12.4 | 11.5 |
| intermediate | 3,115 | 6.4 | 6.0 |
| bright | 39,189 | 80.5 | 75.0 |
| GFP- | 1,485 | 3.0 | 2.8 |
