## Supplementary material for "Mechanisms regulating reactivation pathways of *Toxoplasma gondii* as revealed by bradyzoite transgenesis": Protocol S3

### Protocol S3: Cloning and PCR screening ME49EW transgenic tissue cysts.

Part I. Identifying and infecting mice with single cysts from brain homogenates

1. Prepare and harvest 30-40 d.p.i. mouse brain cortex.

2. **Prepare cortex homogenates and cyst counts.** Briefly, homogenize brain tissue that has been incubated in 3 ml of 1xPBS overnight at 4$^{o}$C by sequential passage through 18g, 20g, and 22 g needles. Count cysts in a 30 µl homogenate spread on slides—use 24X60 coverslips. Cyst counts per slide x 100 estimates the total cysts in the homogenate. (for additional details for preparing homogenates refer to Protocols S1: Cyst and bradyzoite purification.)

3. **Prepare optical plates.** Dilute the homogenate or an aliquot to 10 cysts/ml in 1xPBS. You will need 10 ml of diluted homogenate per optical 96 well plate (SPL Black Plate #33396 or equivalent). Plate 0.1ml per well using a multichannel pipettor being careful to repeat the mixing the homogenate with the pipettor to prevent cysts from settling.

4. **Visually screen for single cysts**. Using suitable microscope equipment, score each well for the presence and number of cysts and GFP fluorescence, if applicable. Record the results on a 96 well map. We use a phase/fluorescent Nikon TS100 Eclipse inverted microscope equipped with 4x, 10x, 20x, and 40x objectives. Alternatively, wells with single cysts can be identified using a TC lab inverted microscope followed by GFP scoring on an inverted fluorescent microscope used for cell image capture. A third alternative is a cell imaging plate reader such as the BioTek Cytation 1.

5. **Infecting mice.** Using a P200 pipette set to ~105 µl, take up the contents of a well containing a single cyst, rinse the edges of the well with this volume, then transfer it into a 1.7 ml microcentrifuge tube. Add 100µl of 1X PBS to the tube (TL vol. 0.2ml). For expected transgenic frequencies 30-50%, infect 10 mice with single cysts. Keep the cysts cool until mouse injections. Using an insulin syringe, take up the entire volume of one tube and inject each mouse i.p.

**Instructions for using a BioTek Cytation 1.**

1. After marking the single cyst wells on a map, open the program “Gen5 3.09” from the desktop computer. Select “Imager Manual Mode”-> “Capture Now” -> “96 Well Optical”.
2. Insert the plate into the imager, choose the well you would like to analyze, and select “Bright Field” from the drop down box in the upper left hand corner.
3. Using the toggle and arrows on the upper left side of the screen locate the cysts according to the map you made above.
4. Change the objective between 4X and 10X using the drop down box in the upper left hand corner if needed.
5. Once a cyst has been located, use the drop down box in the upper left hand corner to change image view from “Bright Field” to “GFP” to determine whether or not the cyst is GFP positive.

Part II. PCR screening for knockout or knock-in transgenic cysts in 96 well plates

**Lysing cysts**.

1. To each well containing cysts (single or more), add 2 µl of Proteinase K (20mg/mL concentration) to each well and place the 96 well plate in the 37℃ incubator for at least 1 hour.

2. Transfer the entire contents of the well into PCR tubes and place them in the PCR machine. Run the following program:

   Incubate at 50$^{o}$C for 60 mins

   Incubate for 95$^{o}$C for 15 minutes

   Hold at 4$^{o}$C.

3. Spin down the contents of the PCR tubes in a microfuge at a speed of 3800 RPM for 15 min.

**Precipitate nucleic acids.**

4. Transfer 90 µl of the supernatant (or as much as possible if there is less than 90 µL) to a 1.7 mL microcentrifuge tube without disturbing the pellet.

5. Add 1/10 volume of 7.5M ammonium acetate, mix, and add 2.5 volumes of cold 100% ethanol. Incubate on ice for an hour.

6. Pellet the DNA in a microfuge at max speed for 30 min. Carefully remove and discard the supernatant, avoiding the pellet (Note: pellet may not be visible)

7. Resuspend the pellet in 150 µl of 70% ethanol and spin down at maximum speed for 30 min.

8. Carefully remove and discard the supernatant, spin again at maximum speed for 5 minutes and carefully remove any excess supernatant.

9. Dry pellet by leaving the tubes open for at least 15 minutes on the bench (or in a fume hood if you have one).

10. Resuspend the pellets in 20 µl of TE Buffer. Use 4 µl in a suitable PCR reaction (20 µl TL vol).

**Note:** To analyze gene knockouts (double crossovers), we design right and left arm primer pairs (inside/outside design) and an inside primer pair designed to amplify a segment of the original gene. Cysts with the correct knockout phenotype, will PCR amplify correct size right and left arm fragments and fail to amplify an inside fragment. For knock-in transgenics such as epitope tagging (single crossover), we screen with an inside/outside primer pair that confirms the correct 3'-end single plasmid insertion. Primers are typically 30-35bp with Tm of 60$^{o}$C or higher and the PCR reaction is 35 cycles (Phusion polymerase). Shorter primers often produce spurious bands due to the crude nucleic acid template used here.

The primer designs for the HXGPRT and AP2IX-9 knockout PCR screens are below:

| Purpose | primer name | Gene | Orientation | Sequence (5' to 3') |
|---|---|---|---|---|
| IX-9 left arm | CH_IX-9_5'F(717) | AP2IX-9 | Forward | GTCATGTGGCATTTCAGTTATGTGTAACATGGTCGAAA |
| | CH_5'R2_gra(662) | pDEST_gfp_DHFR | Reverse | gatacaaggacgcagaaaggaacaaacaccgt |
| IX-9 right arm | CH_DHFR_3'F(656) | DHFR | Forward | GACGAATCCAGATGGAGATGGCT |
| | **CH_IX-9_3'R (291)** | AP2IX-9 | Reversed | CCCGCAGAAAAAAGCCTTATCTTAGG |
| AP2IX-9 inside | CH_IX-9_F2(719) | AP2IX-9 | Forward | GAAAGAAAACTTCGGATGTTGTCAGCGGACTATGC |
| | CH_IX-9_R2(720) | AP2IX-9 | Reverse | GCGAGTCACATGCAGAAAGCAACGAGAATC |

| | | | | |
|---|---|---|---|---|
| HXGPRT Inside | CH_IN_HX_F2(666) | HXGPRT | Forward | atcagaaaaaataagataatggctcctgttctgagtgtttacctgc |
| | CH_IN_HX_R2(667) | HXGPRT | Reverse | CAGAACGTGCTTGTCGCGAAAGATTGACAAG |
| HXGPRT left arm | CH_HX_5'F(664) | HXGPRT | Forward | AAAATTTTTTACAGAAACACCTGGAAATTGATTGCTTCCGCG |
| | CH_5'R2_gra(662) | pDEST_gfp_DHFR | Reverse | gatacaaggacgcagaaaggaacaaacaccgt |
| HXGPRT right arm | CH_DHFR3'F2(663) | pDEST_gfp_DHFR | Forward | tcgccttgttttctttttgtcaatcagaacgaaagcga |
| | CH_HX_3'R2(665) | HXGPRT | Reverse | TGAATAAAATTTCAAATCGCAGAAGTGGATCCGAGAGGC |
