## Supplementary material for "Mechanisms regulating reactivation pathways of *Toxoplasma gondii* as revealed by bradyzoite transgenesis": Protocol S4

**Protocol S4: Cryopreservation of tissue cysts in brain tissue homogenates:**

- Harvest the brain cortex from mice using Protocol S1 (cortex in 15ml conical tube containing 3 ml of 1XPBS stored overnight at 4$^{o}$C). Homogenize sequentially in the conical tube using 18g, then 20g, then 22g needles.
- Freezing whole or portions of brain homogenates
  - Spin down the brain homogenate at 1,000xRPM 5min, remove supernatant (a whole brain will yield 0.5ml - 1 ml pellet volume).
  - Resuspend in freeze media using a ratio of 0.5ml pellet to at least 1ml of freezing media (total volume 1.5 ml).
  - If freezing portions of the brain homogenate in multiple vials, which is especially useful when the homogenate has high cyst numbers (>700 cysts/ml). Spin down the aliquots of homogenate 1000xRPM 5 min, remove supernatants.
  - Again resuspend the pellet using the minimum ratio of 0.5ml pellet volume to 1.0ml freezing media ratio. Higher volumes of freezing media are fine. Aliquot the suspensions into the number of vials desired.
- Incubate the homogenate in freezing media for 15-45 min RT (during this time label tubes)
- Place in Styrofoam box in -80$^{o}$C for 4 h (or o/n). Move to cryo box in -80$^{o}$C freezer
- Freeze Media 1ml recipe:
  - 3-4 drops of Pen/Strep or Penn/Strep/Amphotericin
  - 150μl DMSO
  - 850μl FBS

The major use of frozen cyst stocks is to start up a strain in mice. We are also using frozen homogenates for cloning individual cysts. To thaw homogenate with tissue cysts for mice infections:

- Set up a 15ml conical tube with 5ml of warmed 1x PBS
- Remove cryo tube from -80$^{o}$C freezer and place in 37$^{o}$C water bath
- Quickly thaw (spray outside of tube with 70% alcohol and wipe down before opening) and transfer the homogenate to the 15ml conical tube, gently mix by inversion.
- Centrifuge for 10min at 1500 rpm
- Remove all but 500μl of the PBS (discard), add 3 ml of fresh warm PBS and slowly resuspend.
- Count cysts in a 30μl sample using standard light microscope methods. and dilute as needed for mouse infections.

Notes: Slow cooling in a Styrofoam container or the equivalent in the -80$^{o}$C freezer is important. Tissue cysts in homogenates can be stored in liquid N2 after freezing at -80$^{o}$C. We often see a cyst number drop during the freezing process and then cyst numbers stabilize The Table below shows representative thawed cyst numbers frozen for up to 8 months in a -80oC freezer or in liquid nitrogen.

| | Time frozen | | |
|---|---|---|---|
| Sample type | 2 months | 6 months | 8 months |
| | thawed cyst numbers | | |
| -80$^{o}$C sample 1 | 2058 | 1813 | 1435 |
| -80$^{o}$C sample 2 | 2720 | 1963 | 1517 |
| liquid N2 sample 1 | 1360 | 1533 | 1789 |
| liquid N2 sample 2 | 1430 | 953 | 972 |
